# Structural basis for breadth development in a HIV-1 neutralizing antibody

**DOI:** 10.1101/2022.09.14.507935

**Authors:** Rory Henderson, Ye Zhou, Victoria Stalls, Kevin Wiehe, Kevin O. Saunders, Kshitij Wagh, Kara Anasti, Maggie Barr, Robert Parks, S. Munir Alam, Bette Korber, Barton F. Haynes, Alberto Bartesaghi, Priyamvada Acharya

## Abstract

Antibody affinity maturation enables adaptive immune responses to a wide range of pathogens. In some individuals broadly neutralizing antibodies develop to recognize rapidly mutating pathogens with extensive sequence diversity. Vaccine design for pathogens such as HIV-1 and influenza have therefore focused on recapitulating the natural affinity maturation process. Here, we determined structures of antibodies in complex with HIV-1 Envelope for all observed members and ancestral states of a broadly neutralizing HIV-1 antibody clonal B cell lineage. These structures track the development of neutralization breadth from the unmutated common ancestor and define affinity maturation at high spatial resolution. By elucidating contacts mediated by key mutations at different stages of antibody development we have identified sites on the epitope-paratope interface that are the focus of affinity optimization. Thus, our results identify bottlenecks on the path to natural affinity maturation and reveal solutions for these that will inform immunogen design aimed at eliciting a broadly neutralizing immune response by vaccination.

**Summary:** Somatic hypermutation drives affinity maturation of germline-encoded antibodies leading to the development of their pathogen neutralization function^1^. Rational vaccine design efforts that aim to recapitulate affinity maturation rely on information from antibodies elicited and matured during natural infection. High-throughput next generation sequencing and methods for tracing antibody development have allowed close monitoring of the antibody maturation process. Since maturation involves both affinity-enhancing and affinity-independent diversification, the precise effect of each observed mutation, their role in enhancing affinity to antigens, and the order in which the mutations accumulated are often unclear. These gaps in knowledge most acutely hinder efforts directed at difficult targets such as pan-HIV, pan-Influenza, and pan-Coronavirus vaccines. In HIV-1 infection, antibody maturation over several years is required to achieve neutralization breadth. Here, we determined structures of antibodies in complex with HIV-1 Envelope trimers for all observed members and ancestral states of a broadly neutralizing HIV-1 antibody clone to examine affinity maturation as neutralization breadth developed from the unmutated common ancestor. Structural determination of epitope-paratope interfaces revealed details of the contacts evolving over a timescale of several years. Structures along different branches of the clonal lineage elucidated differences in the branch that led to broad neutralization versus off-track paths that culminated in sub-optimal neutralization breadth. We further determined structures of the evolving Envelope revealing details of the virus-antibody co-evolution through visualization of how the virus constructs barriers to evade antibody-mediated neutralization and the mechanisms by which the developing antibody clone circumvents these barriers. Together, our structures provide a detailed time-resolved imagery of the affinity maturation process through atomic level descriptions of virus-antibody co-evolution leading to broad HIV neutralization. While the findings from our studies have direct relevance to HIV-1, the principles of affinity optimization and breadth development elucidated in our study should have broad relevance to other pathogens.

## Introduction

Antibody affinity maturation from germline-encoded heavy and light chain immunoglobulin (Ig) genes involves iterative rounds of somatic hypermutation and selection followed by B cell expansion and differentiation, ultimately leading to pathogen neutralization^1^. The advent of high-throughput next generation sequencing and methods for tracing antibody development has allowed close monitoring of the affinity maturation process^2,3^. Extensive structure-based studies of antibodies alone or of antibodies in contact with cognate antigen have revealed that affinity gains involve acquisition of mutations that improve antibody-antigen contacts, shape complementarity, paratope rigidity, and antibody conformation^2,4–8^. Maturation also involves affinity-independent diversification, which, together, leads to a broad range of potential solutions to the affinity optimization problem^4,9^. Antibodies with neutralization breadth can effectively bind to target antigen despite sequence variability and are a major vaccine design target for pathogens such as HIV-1, influenza, and SARS-CoV-2 and other human coronaviruses^10–12^.

While influenza and SARS-infections are usually cleared within a relatively short time after infection, HIV-1 is a chronic, genome-integrating infection and antibody maturation resulting in broadly neutralizing antibodies (bnAbs) only occurs over multiple years. Defining antibody maturation pathways to strategically inform immunogen design and enable acceleration of this process through vaccination is a primary goal of current HIV-1 vaccine efforts^13^. Broadly neutralizing antibodies have been isolated from people living with HIV (PLWH) and provide the basis for many vaccine strategies aiming to induce bnAbs^14^. Development of HIV-1 directed bnAbs typically requires a defined set of specifically positioned mutations in heavy/light chain Ig pairs to occur in the antibody clone, presenting a formidable challenge in immunogen design aimed at recapitulating bnAb development. This is exacerbated by HIV-1 bnAb features that are uncommon including extensive somatic mutations, long heavy chain third complementarity determining regions (HCDR3s), Ig insertions and deletions, restricted germline gene usage, and enrichment for low probability mutations^15–17^. A detailed understanding of the affinity maturation steps leading to bnAb development, including discrimination among acquired mutations that lead to the desired bnAb response versus those that culminate in less productive off-target responses, is therefore essential to identify the determinants of affinity matured bnAb B cell receptor (BCR) selection.

Studying the coevolution of virus and antibody clones during HIV infection informs vaccine design by defining the HIV-1 Envelope (Env) variants that evolve during bnAb development, thus providing a blueprint for iterative immunogen design^18–21^. The targets for HIV-1 bnAbs are conserved epitopes on the Env protein^22–25^. The HIV-1 Env is a trimer of gp120-gp41 heterodimers that are heavily shielded from the host immune systems by N-linked glycosylation and further protected from neutralization by conformational masking and considerable sequence variability.

A glycosylated region near the base of the third variable loop (V3) of HIV-1 Env forms a supersite of vulnerability that is targeted by antibodies originating from diverse germline genes in multiple HIV-1 infected individuals^26^. The development of a broadly neutralizing V3-glycan antibody was studied in an African male living with AIDS from Malawi (CH848), who was followed from the time of infection up to 5 years after transmission^21^. In the CH848 individual, the early appearance of two autologous “cooperating” neutralizing B cell clones (DH272 and DH475) led to the selection of viral escape variants, that in turn stimulated the DH270 clone that further developed to acquire potent neutralization breadth^21^. DH270 antibodies were detected in the CH848 individual at week 186 and coincided with the appearance of Env variants with a shortened variable loop 1 (V1)^21^. The DH270 unmutated common ancestor antibody (DH270.UCA), representing the naïve B cell progenitor of the DH270 clone, does not neutralize heterologous HIV-1, although a single amino acid change at position 57 of the heavy chain that substituted a glycine for an arginine (G57R) resulted in heterologous HIV-1 neutralization, albeit with limited breadth. As the DH270 clone affinity matured and antibodies accumulated mutations in their heavy and light chain variable regions (V_H_ and V_L_), breadth and potency of heterologous neutralization improved. The early DH270 ancestral intermediate antibodies were able to neutralize heterologous viruses with short V1 loops. The DH270 B cell clonal lineage evolved to gain capacity to neutralize viruses with longer V1 loops, although with reduced potency^21^. Similar inverse correlation between potency and V1 length was observed for other V3-glycan bnAbs 10-1074, PGT121, and PGT128^27,28^. The rich dataset of co-evolving antibodies and Envs available for the DH270 clone^21^ presented an opportunity to study structural aspects of affinity maturation and virus coevolution at an unprecedented level of detail and to define how this information will inform vaccine design aimed at eliciting bnAbs targeting the V3-glycan supersite.

Three key discoveries have informed our current understanding of the DH270 clone. First, though somatic hypermutation is a stochastic process, hot spots and cold spots in the antibody sequence exist, leading to differences in the probabilities of which antibody sites are mutated and of the residues these are mutated into. HIV-1 bnAbs, including those in the DH270 clone, are enriched for key, low probability mutations^17^. Second, of the forty-two mutations in the most broad and potent member in this clone, only twelve are needed to reach ninety percent of its breadth, allowing us to pinpoint the most critical regions of the antibody that need to be optimized^29^. Finally, key for considering affinity maturation in the context of vaccine development, we have shown that rationally designed immunogens can expand DH270 bnAb V3-glycan precursors ^22^.

Thus, to understand antibody evolution over time at a structural level, here we used cryo-EM to visualize the interactions of the complete DH270 clonal tree with the co-evolving virus Env. We define the defenses mounted by the virus to shield its V3-glycan epitope during infection and demonstrate how the DH270 clone developed to first engage Env and then matured to effectively circumvent these barriers to achieve neutralization breadth. Our results show that clone development involved sequential solutions to affinity gain at specific antibody-antigen contact sites. At each stage of development, the acquisition of a solution was followed by splitting of the clone into distinct sub-clones leading to differential gains in affinity and neutralization breadth. These results showed that mutations acquired early in the clone determined the fate of downstream DH270 clonal affinity maturation. Antibody maturation involved staged, site specific optimization of epitope-paratope contacts with the site of focus shifting as evasive mutations accumulated on the antigenic surface. These findings support a vaccine development pipeline that sequentially optimizes specific antibody properties at spatially distinct sites.

## Results

### Structural determination of the Env bound DH270 clone antibody Fabs

The structures of several DH270 antigen binding fragments (Fabs) have been previously determined, including inferred DH270 UCAs, as well as the mature DH270.3, DH270.5, and DH270.6 antibodies that were isolated from the CH848 individual. Additionally, structures have been determined of a DH270.1 single-chain variable fragment (scFv), a Mannose associated DH270.3, a V3-glycan peptide associated DH270.6 scFv, and soluble Env ectodomain bound DH270 UCA and DH270.6 Fabs^21,22,30^. While the V3-glycan associated DH270.6 structure clarified the role of individual residues critical for neutralization breadth, it did not reveal the structural pathway underlying the evolution of this breadth, nor show how breadth and potency were affected by mutations in different DH270 clades and subclades. We therefore sought to determine the impact of mutations at each stage of maturation by determining cryo-EM structures of HIV-1 Env associated DH270 clonal member Fabs from each antibody ancestral intermediate, and all mature antibody forms (Figure 1 and Supplemental Figures 1 and 2). Unless otherwise noted, a soluble SOSIP trimer derived from a virus isolated from the CH848 individual at day 949 (referred to here as the d949 Env) after transmission was used for preparing complexes with DH270 clone Fabs for structural studies. Map resolutions ranged from 5.3-3.3 Å with local resolutions at the Fab-Env interface in the range of 5-3 Å (Supplemental Figures 2–4, Table 1).

**Figure 1.**
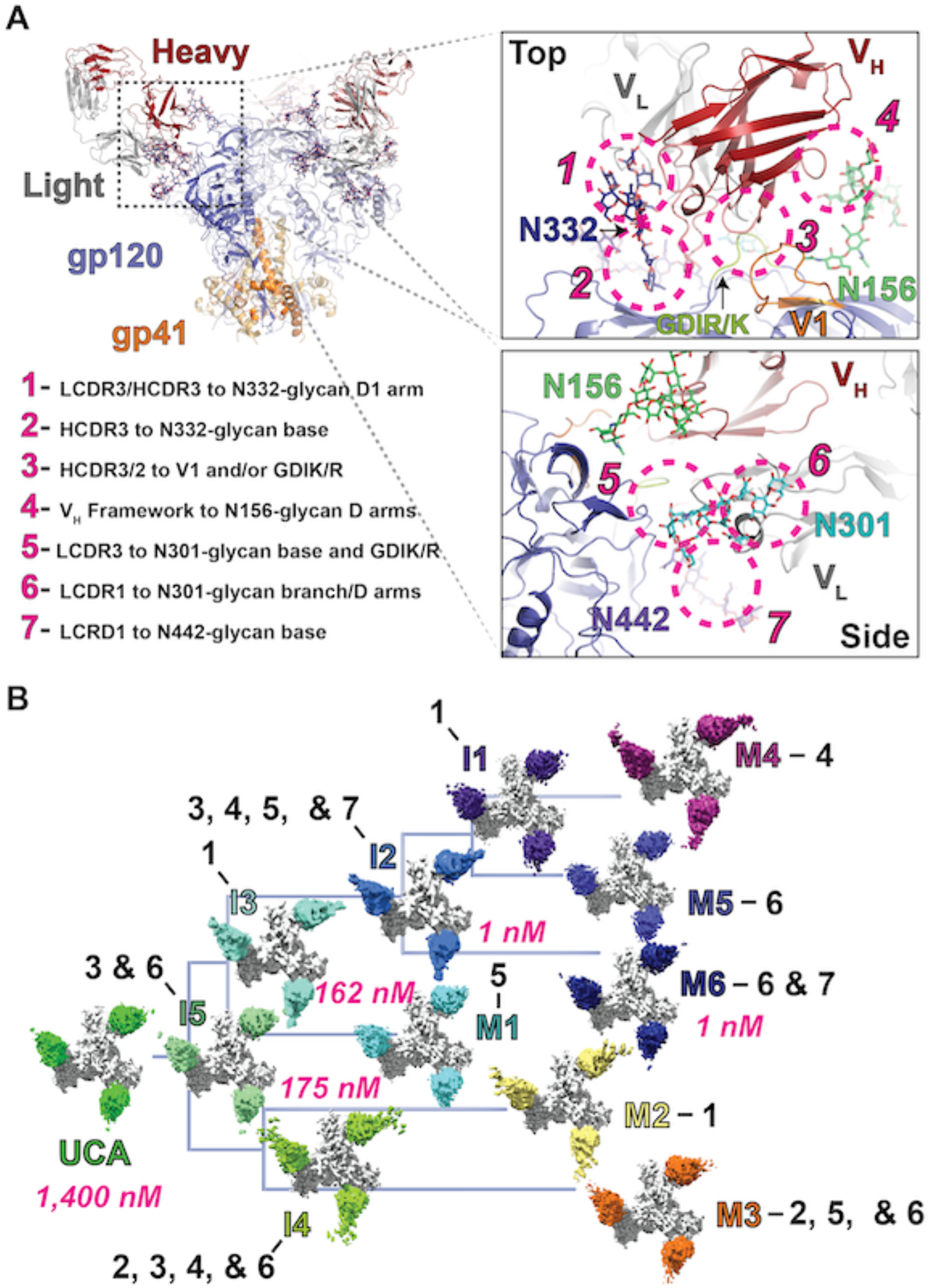
The V3-glycan targeting DH270 broadly neutralizing antibody clone. **A)** (upper left) A DH270 antibody Fab bound HIV-1 Envelope (Env) ectodomain highlighting the gp120 (blue) and gp41 (orange) subunits. Antibody Fab heavy and light chains are colored red and grey, respectively, with epitope glycans depicted in stick representation. (right) Zoomed-in, top and side views of the paratope-epitope contacts. Dashed circles indicate distinct contact regions. (lower left) Number indicators identify contact sites highlighted by the dashed circles in the panels to the right. **B)** The inferred DH270 clone phylogenetic tree depicting cryo-EM maps determined for each clonal member Fab bound to the CH848 d949 trimer (white). The clonal member names are colored according to the Fab color of each map. Black numbering indicates locations of consequential mutations in each antibody according to sites listed in (*A*). Affinities for antibodies on the path leading to the most broad and potent mature form, M6, are indicated in pink. Intermediate clonal members are denoted with an I while mature antibodies are denoted with an M.

**Table 1.** Cryo-EM Data Collection and Refinement Statistics.

|  | (DH270.UCA) | (DH270.UCA.G57R-CH848.10.17DT) | (DH270.I5.6-CH848.10.17) | (DH270.I4.6-CH848.10.17) | (DH270.I3-CH848.10.17) | (DH270.I2-CH848.10.17) | (DH270.I1.6-CH848.10.17) | (DH270.1-CH848.10.17) | (DH270.2-CH848.10.17) | (DH270.3-CH848.10.17) |
| --- | --- | --- | --- | --- | --- | --- | --- | --- | --- | --- |
| **Data Collection** |  |  |  |  |  |  |  |  |  |  |
| Microscope |  |  |  |  |  | FEI Titan Krios |  |  |  |  |
| Detector |  |  |  |  |  | Gatan K3 |  |  |  |  |
| Magnification |  |  |  |  |  | 105000 |  |  |  |  |
| Voltage (kV) |  |  |  |  |  | 300 |  |  |  |  |
| Electron dose (e-/Å<sup>2</sup>) | 71 | 69 | 53 | 60 | 60 | 60 | 60 | 60 | 60 | 60 |
| Pixel Size (Å) | 1.06 | 1.066 | 1.066 | 1.08 | 1.066 | 1.066 | 1.04 | 1.066 | 1.066 | 1.066 |
| Defocus | ~0.8- | ~0.7-4.1 | ~0.5-4.1 | ~0.5-3.1 | ~0.5-3.5 | ~0.5-3.5 | ~0.5-4 | ~1.5-3.5 | ~0.5-4 | ~0.5-3.5 |
| Range (µm) | 4.3 |  |  |  |  |  |  |  |  |  |
| Reconstruction software |  |  |  |  |  | cryoSPARC |  |  |  |  |
| Symmetry | C3 | C3 | C3 | C3 | C3 | C3 | C3 | C3 | C3 | C3 |
| Particles | 120981 | 208426 | 101922 | 338,934 | 67110 | 39612 | 65522 | 125676 | 24465 | 76274 |
| Box size (pix) | 384 | 384 | 384 | 300 | 384 | 384 | 384 | 384 | 384 | 384 |
| Resolution (Å) | 3.8 | 3.3 | 3.9 | 4.2 | 4.3 | 3.5 | 3.9 | 3.6 | 4.1 | 3.4 |
| (FSC0.143)<sup>\$</sup> | | | | | | | | | | |
| **Coordinate Refinement (Phenix)<sup>#</sup>** |  |  |  |  |  |  |  |  |  |  |
| Protein residues | 2496 | 2490 | 2490 | 2490 | 2490 | 2490 | 2490 | 2490 | 2490 | 2490 |
| **R.M.S. deviations** |  |  |  |  |  |  |  |  |  |  |
| Bond lengths (Å) | 0.006 | 0.006 | 0.006 | 0.007 | 0.006 | 0.006 | 0.006 | 0.013 | 0.006 | 0.006 |
| Bond angles (°) | 0.908 | 0.892 | 0.909 | 0.987 | 0.898 | 0.990 | 0.885 | 1.159 | 0.926 | 0.933 |
| **Validation** |  |  |  |  |  |  |  |  |  |  |
| Molprobrity score | 1.42 | 1.48 | 1.39 | 1.48 | 1.57 | 1.56 | 1.42 | 1.44 | 1.54 | 1.52 |
| Clash score | 2.15 | 2.86 | 2.30 | 2.56 | 3.06 | 3.95 | 2.59 | 2.91 | 2.54 | 2.98 |
| Favored rotamers (%) | 99.77 | 99.53 | 99.72 | 99.63 | 99.67 | 99.21 | 99.86 | 99.72 | 99.53 | 99.68 |
| EMRinger Score | 2.12 | 2.06 | 2.28 | 1.81 | 2.73 | 2.53 | 2.61 | 2.23 | 1.39 | 1.63 |
| **Ramachandran plot** |  |  |  |  |  |  |  |  |  |  |
| Favored regions (%) | 93.41 | 94.04 | 94.4 | 93.11 | 92.34 | 94.48 | 94.44 | 94.77 | 91.65 | 93.31 |
| Disallowed regions (%) | 0.2 | 0.36 | 0.12 | 0.24 | 0.24 | 0.16 | 0.36 | 0.32 | 0.49 | 0.28 |

<sup>\$</sup> Resolutions are reported according to the FSC 0.143 gold-standard criterion
<sup>@</sup> A highly heterogeneous dataset with many 3D classes. The top two classes were refined with C1 symmetry.
<sup>#</sup> Statistics are reported for the protein residues within the complex excluding the antibody constant domains

**Table 2.** Cryo-EM Data Collection and Refinement Statistics.

|  | (DH270.4<br>-<br>CH848.1<br>0.17) | (DH270.5-CH848.10.1<br>7) | (DH270.6-CH848.10.17) | (VRC01-CH848.0526.<br>25) | (VRC01-CH848.10.17<br>) | (DH270.6-CH848.0526.<br>25) | (VRC01-CH848.0836.<br>10) | (VRC01-CH848.0358.<br>80) |
| --- | --- | --- | --- | --- | --- | --- | --- | --- |
| **Data Collection** |  |  |  |  |  |  |  |  |
| Microscope |  |  |  |  | FEI Titan Krios |  |  |  |
| Detector |  |  |  |  | Gatan K3 |  |  |  |
| Magnification |  |  |  |  | 105000 |  |  |  |
| Voltage (kV) |  |  |  |  | 300 |  |  |  |
| Electron dose (e<sup>-</sup><br>/Å<sup>2</sup>) | 60 | 60 | 53.15 | 69.98 | 60 | 60 | 60 | 60 |
| Pixel Size (Å) | 1.066 | 1.066 | 1.066 | 1.066 | 1.066 | 1.066 | 1.066 | 1.066 |
| Defocus Range<br>(µm) | ~0.8-4.3 | ~0.7-4.1 | ~0.5-4.1 | ~0.5-3.1 | ~0.5-3.5 | ~0.5-3.5 | ~0.5-3.5 | ~0.5-3.5 |
| Reconstruction |  |  |  |  | cryoSPARC |  |  |  |
| Software |  |  |  |  |  |  |  |  |
| Symmetry | C3 | C3 | C3 | C3 | C3 | C3 | C3 | C3 |
| Particles | 107890 | 76211 | 106823 | 252873 | 35839 | 119542 | 28746 | 46336 |
| Box size (pix) | 384 | 384 | 384 | 384 | 384 | 384 | 384 | 384 |
| Resolution (Å) | 3.3 | 4.0 | 3.82 | 4.0 | 4.5 | 3.9 | 4.6 | 4.9 |
| (FSC0.143)<sup>\$</sup> | | | | | | | | |
| **Coordinate Refinement (Phenix)<sup>#</sup>** |  |  |  |  |  |  |  |  |
| Protein residues | 2490 | 2496 | 2490 | 2514 | 2457 | 2574 | 2505 | 2523 |
| **R.m.s. deviations** |  |  |  |  |  |  |  |  |
| Bond lengths<br>(Å) | 0.006 | 0.008 | 0.006 | 0.004 | 0.005 | 0.006 | 0.004 | 0.006 |
| Bond angles (°) | 0.915 | 1.100 | 1.055 | 0.845 | 0.894 | 1.041 | 0.949 | 0.997 |
| **Validation** |  |  |  |  |  |  |  |  |
| Molprobrity score | 1.57 | 1.71 | 1.71 | 1.53 | 1.80 | 1.76 | 1.71 | 1.90 |
| Clash score | 3.13 | 4.02 | 4.37 | 2.96 | 5.01 | 4.26 | 4.65 | 6.60 |
| Favored rotamers<br>(%) | 99.72 | 99.68 | 99.30 | 99.77 | 99.81 | 99.73 | 99.59 | 100 |
| EMRinger Score | 1.95 | 0.36 | 1.74 | 1.66 | 1.65 | 1.49 | 1.31 | 0.56 |
| **Ramachandran plot** |  |  |  |  |  |  |  |  |
| Favored<br>regions (%) | 92.46 | 90.94 | 91.93 | 93.14 | 90.56 | 89.78 | 92.51 | 90.15 |
| Disallowed<br>regions (%) | 0.32 | 0.44 | 0.41 | 0.24 | 0.41 | 0.35 | 0.12 | 0.24 |

<sup>\$</sup> Resolutions are reported according to the FSC 0.143 gold-standard criterion
<sup>@</sup> A highly heterogeneous dataset with many 3D classes. The top two classes were refined with C1 symmetry.
<sup>#</sup> Statistics are reported for the protein residues within the complex excluding the antibody constant domains

Mapping the mutations of the DH270 clone on the cryo-EM structures revealed clustered sites that are sequentially mutated along the B cell clonal tree (Figure 1A). These clusters mapped to seven distinct sites of antibody/Env contact: (1) interaction of the distal D1 arm of the Env N332-glycan with the cleft formed between the Fab variable heavy and light chain segments (V_H_/V_L_) involving the light-chain complementarity determining region two (LCDR2) and heavy-chain complementarity determining region three (HCDR3), (2) interaction of the N332-glycan GlcNac base with the antibody HCDR3 region, (3) contact of the Env V1 loop, and of the conserved GDIR/K motif in the V3 loop with the Fab HCDR3 and HCDR2 regions, (4) interaction of the D arms of the Env N156-glycan with the antibody framework region, (5) interaction of the Env N301-glycan base GlcNac-1 and residues around the V3 GDIR/K motif with the Fab LCDR3 region, (6) contact of the Env N301-glycan branch point and D arms with the Fab LCDR1/V_L_ N-terminus site, and (7) contact of the Env N442-glycan with Fab LCDR1 region (Figure 1A). Together, these comprise the entirety of the interactive surface available to the DH270 clone antibodies.

The cryo-EM reconstructions were well-resolved at the Ab/Env interface. Clear patterns of mutations emerged along the DH270 clonal tree at the Env interactive sites described above that were suggestive of the DH270 clone evolving to resolve specific, sequential structural bottlenecks on the path of affinity maturation (Figure 1B). The first intermediate, I5, acquired mutations that strengthened interactions at clusters 3 and 6 by improving protein-protein contacts as well as the interactions of the antibody with the N301 glycan, resulting in ∼8-fold improved binding affinity to the d949 Env relative to the UCA. After intermediate I5, the clone splits into two distinct paths along a clade defined by the I3 intermediate and a clade defined by the I4 intermediate. This consequential split leads to the acquisition of greater neutralization breadth and potency in the I3 clade, leading to the DH270.6 bnAb (Figure 1B)^21^. Both clades acquired mutations around the N332-glycan interaction site but at differing positions. The I3 intermediate improves interaction with the distal N332-glycan D1 arm (cluster 1) while the I4 intermediate improves interaction with the N332-glycan GlcNac base (cluster 2). Each intermediate then splits into two subclades, revealing a conserved Env target for contact optimization by three of the four mature/intermediate antibodies. These include the I4 clade mature antibody DH270.3, I3 clade mature antibody DH270.1, and intermediate antibody I2, which all show several mutations clustered in or adjacent to the antibody LCDR3 region, presumably improving interactions with the Env N301-glycan base (cluster 5). The I4 clade mature antibody DH270.2 acquired mutations focused on cluster 1, improving interactions at the N332-glycan D1 arm in a manner similar to the I3 intermediate. The I2 intermediate acquired additional mutations at clusters 3, 4, and 7, with mature DH270.6, the most broad and potent antibody of the clone, further modifying interactions at cluster 7. A minimally mutated DH270 antibody (minDH270) was designed^29^ where only 12 of the 42 mutations in DH270.6 were needed to reach ∼90% of the breadth and potency of DH270.6. Importantly, the mutations included only those that appeared in I5, I3, and I2. Consistent with this observation, mutations after I2 occurred at more distal sites and their effects were less clear compared to the mutations that appeared in the previous members of the DH270 clone, as described below. The DH270.5 and DH270.6 antibodies acquired several mutations near the LCDR1/V_L_ N-terminus where interactions with the N301-glycan D arms may occur. The cryo-EM reconstructions in this region were poorly resolved, precluding a precise definition of these interactions. Together, these results indicate that maturation of the DH270 clone involved sequential affinity gain at distinct sites, and that specific structural solutions were necessary for affinity gain and development of neutralization breadth. This is exemplified in the dichotomy where acquisition of mutations in the antibody LCDR3 region led to neutralization breadth gain in DH270.1 and in the I2 intermediate, but loss in neutralization breadth in DH270.3.

### UCA to I5: Early mutations in clone development set the stage for downstream maturation

We next studied the specific structural arrangements underlying maturation at each stage, first examining mutations in the I5 intermediate (Figure 2A and B). We previously determined structures for the DH270 UCA and the mature DH270.6 Fabs bound to the CH848 d949 Env timer with V1 loop glycans at positions 133 and 138 removed^22^. In the UCA-bound structure the V1 loop interacts with both the antibody as well as the GDIK motif and adjacent V3 residues, while in the DH270.6-bound Env the V1 loop was displaced from this position. This displacement was attributed to the V_H_ G57R mutation that occurs in the first intermediate^22^. To visualize the V1 loop conformation when not bound to an antibody at the V3 glycan bNAb epitope, we determined the structure of the d949 SOSIP Env in complex with the CD4 binding site antibody VRC01. We found that the conformation of the unbound V1 loop resembled that of the UCA-bound V1 loop (Supplemental Figure 6A). We further determined the structure of DH270.UCA incorporating the V_H_ G57R mutation in complex with the CH848 d949 SOSIP Env (Figure 2C and Supplemental Figure 6B). The V1 loop in the UCA+G57R structure was displaced from its position over the GDIK motif observed in its free and UCA-bound forms. The displacement of the V1 loop in the UCA+G57R structure is accompanied by a marked shift in the Ab orientation, rotating the Fab V_H_/V_L_ toward the space previously occupied by the V1 loop, drawing the antibody closer to the Env (Figure 2C and Supplemental Figure 6C). Comparing the structure of the UCA+G57R-bound complex with the I5-bound complex revealed further shifts in the orientation of the antibody orchestrated by the other acquired mutations in the I5 intermediate. The V_L_ S27Y substitution near the β-Mannose branchpoint of the N301-glycan is the likely culprit, altering the orientation of N301 glycan in the I5-bound complex relative to the UCA+G57R bound complex (Figure 2D). We next quantified these shifts in antibody orientation using vector-based angles between the antibody F_v_ and its gp120 epitope. Centroids for the epitope, the F_v_, and an anchor point in the V_H_ were used to determine the distance between the F_v_ and its epitope and angle (theta), with an additional anchor point in the epitope added to examine rotation (phi) of the antibody relative to the epitope (Figure 2E). Theta and phi describe the antibody angle of approach with reference to the epitope, allowing close inspecting of differences in antibody disposition as mutations accumulate in the clone. The distance between the antibody and the epitope was reduced by ∼0.5 Å in the UCA+G57R bound structure relative to the UCA-bound structure, with further minor reductions observed in I5 associated with the additional I5 mutations (Figure 2E). Comparison of the angle and dihedral dispositions of the UCA and UCA+G57R structures showed a rotation in each of ∼2°. The I5 intermediate showed further rotation about the phi dihedral in the direction of the S27Y mutation without additional changes in the theta rotation angle. Together, these results demonstrate that early mutations in the DH270 clone facilitate improved contacts, shift the position of the antibody relative to the bound Env, and alter Env conformation particularly of the V1 loop and N301 glycan, with the angular and conformational changes further strengthening antibody-Env interactions.

**Figure 2.**
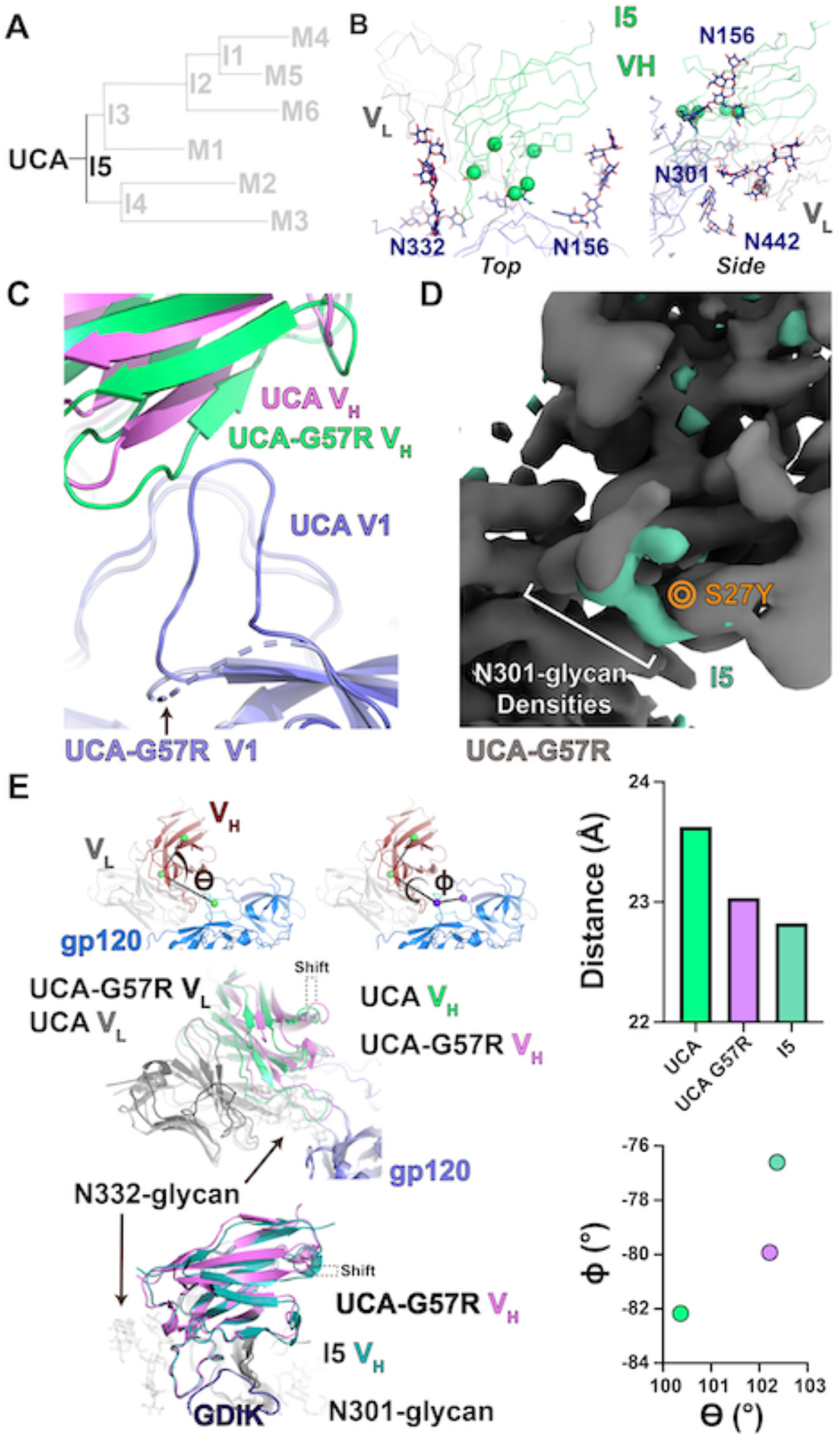
The UCA to I5 transition. **A)** DH270 clonal tree highlighting the UCA and I5 intermediate antibodies. **B)** Top and side views of the I5 contact site. Alpha-carbons (Cα) of heavy and light chain mutation are presented as spheres. **C)** Top view of the UCA and UCA+G57R heavy chain contacts with gp120 highlighting the shifts in the antibody position and in the Env gp120 V1 loop. **D)** Gaussian filtered map of the UCA+G57R Fab bound trimer complex (grey) overlaid with a filtered and subtracted map of the I5 Fab bound trimer complex (green). The location of the S27Y mutation on the maps is indicated in orange. **E)** (upper left) Contact angle theta and dihedral phi with positions of the centroids used for calculations indicated with spheres. (middle left) Alignment (gp120 only) of UCA and UCA+G57R bound state structures highlighting the shift in antibody disposition about the theta angle. (lower left) Alignment (gp120 only) of UCA+G57R and I5 bound state structures highlighting the shift in antibody disposition about the phi dihedral. (upper right) Antibody Fab V_H_/V_L_ centroid distance from the gp120 epitope centroid. (lower right) Theta vs. phi plot for the UCA, UCA+G57R, and I5 structures. The colors of the data points match the respective heavy chain colors in the structures.

### The I5 path split: Mutations in I3 set the path toward broad neutralization

We next examined the early development of two distinct clades in the DH270 clone from the I5 intermediate (Figure 3A and B). The I3 intermediate that occurs after I5 in the DH270.6 clone incorporates heavy chain mutations V11M, R87T, and the improbable R98T mutation, and light chain mutations L48Y and S54N. The cryo-EM reconstruction of the I3 Fab bound Env was determined to a resolution of 4.5 Å. The V1 loop conformational change induced by the V_H_ residue R57 was retained as was the interaction of V_L_ residue Y27 with the N301-glycan (Figure 3B and Supplemental Figure 7A and B). No major differences were observed in the I5 and I3 Fab bound Env structure gp120 and gp41 subunits (RMSDs ∼0.9 Å; Supplemental Figure 7C). As previously described^30^, the V_L_ L48Y and V_H_ R98T mutations introduce new hydrogen bonding contacts, with the Y48 hydroxyl interacting with the N332-glycan, and the release of D115 through the R98T mutation allowing closer interaction of D115 with the N332-glycan (Figure 3C and Supplemental Figure 7D). Additionally, the I3 V_H_ R98T substitution introduced a hydrogen bond between the side chains of residues T98 and the V_H_ Y27 (Supplemental Figure 7E), thus bolstering internal stability of the antibody structure by compensating for the loss of the cation-π interaction of R98 with Y27. The V_H_ R87T and V_L_ S54N mutations are distant from the gp120-interactive region of the antibody. Close examination of the local regions of these mutations suggested potential roles of these substitutions in augmenting internal stability of the antibody. The V_H_ R87T substitution relieves electrostatic strain contributed by three spatially clustered arginines, R67, R85 and R87 (Figure 3E). Notably, a second arginine in this cluster, R85, is mutated to serine at the next step as I3 transitions to I2. Moreover, the T87 side chain makes a hydrogen bond with main chain nitrogen of residue V_H_ D89, thus augmenting stability of local structure. The effect of V_L_ S54N was more subtle with N54 appearing to stabilize interaction with an adjacent beta strand via hydrogen bonding of its side chain with the main chain of light chain residue 66 (Supplemental Figure 7G).

**Figure 3.**
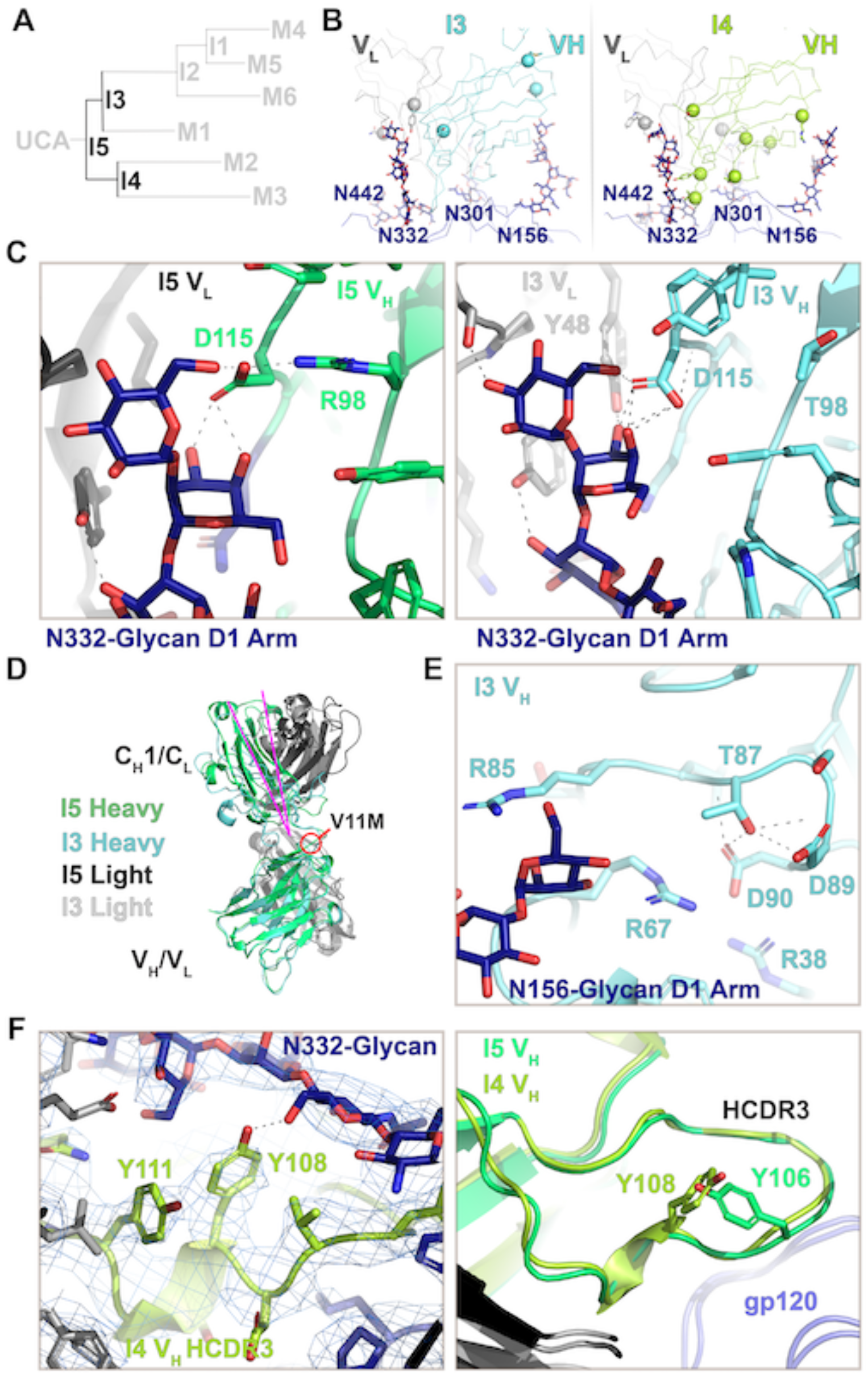
The I5 to I3 and I4 path split. **A)** DH270 clonal tree highlighting the I5, I3, and I4 intermediate antibodies. **B)** Top views of the I3 and I4 contact sites. Alpha-carbons (Cα) of heavy and light chain mutations are presented as spheres. **C)** (left) The N332-glycan D1 arm contact with the I5 V_H_/V_L_ cleft. (right) The N332-glycan D1 arm contact with the I3 V_H_/V_L_ cleft. **D)** Aligned I5 and I3 V_H_ domains highlighting the change in elbow angle (magenta). **E)** Structure of the I3 intermediate at the R87T site near potential N156-glycan contacts. **F)** (left) Map and fit coordinates of the new N332-glycan contact formed at the I4 HCDR3 S108Y mutation site. (right) Aligned I5 and I4 V_H_ domains showing HCDR3 loop arrangements.

As predicted by previous molecular dynamics simulations^6^, the V11M mutation induced a marked shift in the antibody Fab elbow hinge (Figure 3D). Unlike I3, the I4 intermediate did not acquire an elbow mutation and consequently did not show a distinct shift in the elbow region (Supplemental Figure 7H). As in I3, the R57 residue in I4 showed evidence of shifting the position of the V1 loop (Supplemental Figure 7I). However, additional density was observed in this region that is consistent with a GDIK coupled V1 loop and suggestive of multistate behavior. The I4 intermediate acquired a larger number of V_H_ mutations including two in the HCDR3, Y106V and S108Y that essentially shifts the position of the tyrosine from residue 106 to 108. The hydroxyl group in I4 Y108 formed a hydrogen bond with the N332 GlcNac-2 (Figure 3F left). The overall backbone conformation of the HCDR3 region was not altered by these mutations (Figure 3F right). The I4 S84R mutation is in proximity to the distal sugar units of the N156-glycan. Densities for the side chain and this portion of the N156-glycan were, however, not visible in the cryo-EM reconstruction (Supplemental Figure 7J and K). The remaining mutations in the I4 heavy chain at positions G49A, N54T, Q62R, and Y116S play no clearly decipherable role in the interaction with Env or in modifying the antibody structure. The light chain contains two non-paratope mutations, R56W and V100I. No changes were observed around V100I position relative to I5 or I3. Residue R59W of the I4 light chain that is adjacent to the LCDR2 region displayed a modest rearrangement that shifts the position of distal N332-glycan interactive residues (Supplemental Figure 7L). This was not associated with apparent changes in antibody interactions with the glycan though it may confer stabilization of the LCDR2 loop and therefore the glycan interaction. These observations show that at this stage of development, interactions with the N332-glycan were the focus of optimization in both clades of the DH270 clonal tree.

### Maturation after I4: DH270.2 and DH270.3 attempt improvements in N332-glycan interactions or GDIK and N301-glycan interactions

The I4 clade leads to mature antibodies DH270.2 and DH270.3 (Figure 4A and B). DH270.2 showed greater breadth and potency than DH270.3 with each neutralizing ten and fifteen members from a 24-virus heterologous panel, respectively, although both displayed weaker neutralization breadth and potency compared to the I3 clade mature antibodies that neutralize sixteen to seventeen members of the heterologous panel^22^. Alignment of the Env-bound structures revealed distinct shifts in the orientation of each antibody compared to the I4 intermediate (Figure 4C and Supplemental Figure 8A). Inspection of the theta and phi angles indicated DH270.3 had predominantly shifted its rotation about the epitope (Figure 4D). The most notable mutation in DH270.3 was the light chain Y27S reversion. A clear shift in the position of the N301-glycan was observed relative to the I4 intermediate (Figure 4E and Supplemental Figure 8B). Consistent with the observed rotation of the bound antibody in the I5/Env complex relative to the UCA+G57R/Env complex, the Y27S reversion resulted in DH270.3 occupying an orientation matching UCA+G57R (Figure 4F). This reversion of antibody orientation suggests that the S27Y substitution, first acquired in I5, is the driver of the antibody rotation and N301-glycan re-orientation that was observed in going from the UCA to I5.

**Figure 4.**
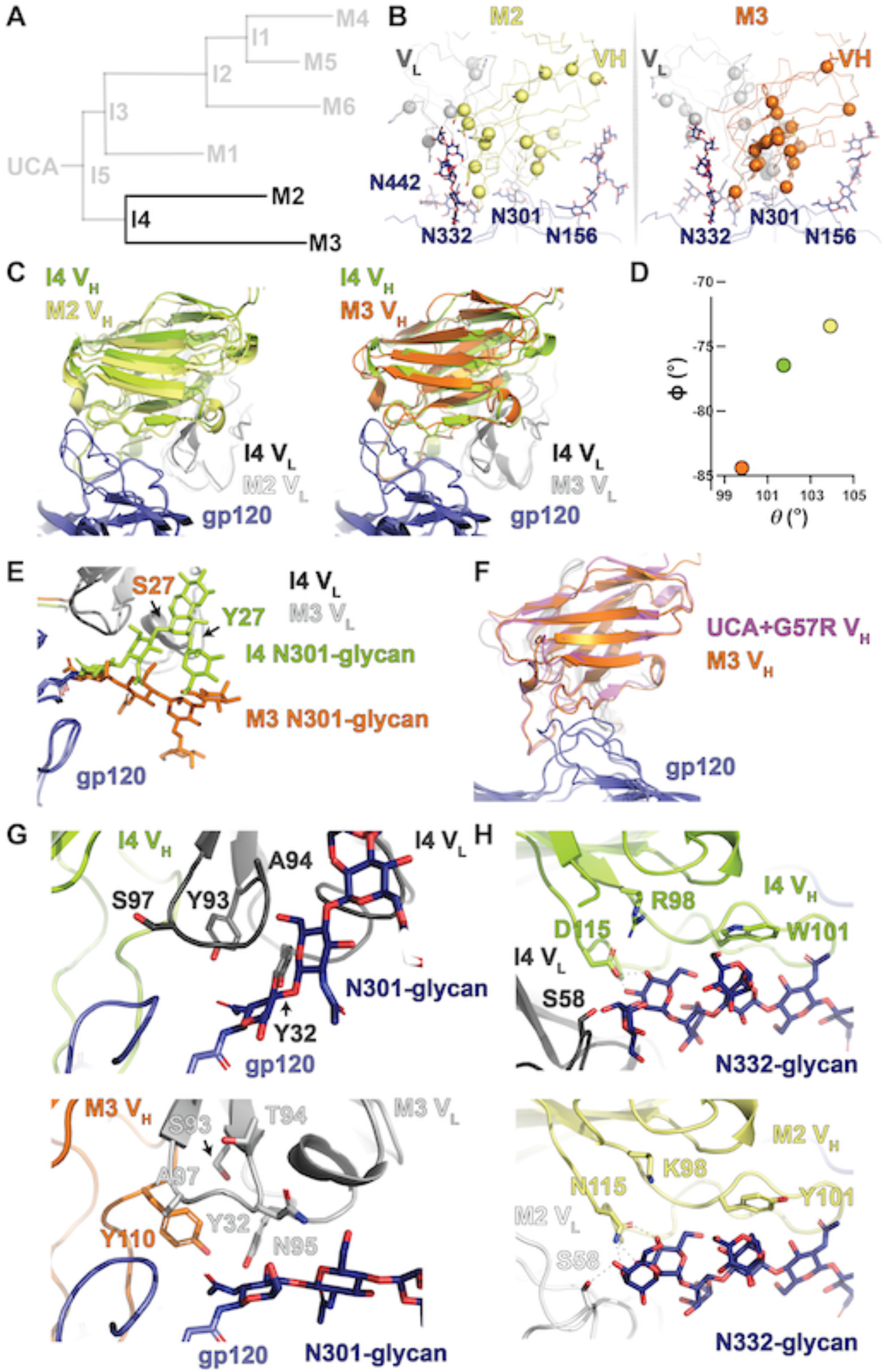
The I4 to mature DH270.2 and DH270.3 transition. **A)** DH270 clonal tree highlighting the I4 intermediate antibody and the mature DH270.2 (M2) and DH270.3 (M3) antibodies. **B)** Top views of the DH270.2 and DH270.3 contact sites. Alpha-carbons (Cα) of heavy and light chain mutations are presented as spheres. **C)** (left) Alignment (gp120 only) of I4 and DH270.2 bound state structures highlighting the shift in antibody disposition (right) Alignment (gp120 only) of I4 and DH270.3 bound state structures highlighting the shift in antibody disposition. Theta vs. phi plot for the I4, DH270.2, and DH270.3 structures. The colors of the data points match the respective heavy chain colors in the structures. Structure overlay of the I4 and DH270.3 N301-glycans showing their change in disposition and the location of the S27 or Y27 residues on the antibody light chain. **F)** Alignment (gp120 only) of UCA+G57R and DH270.3 bound state structures indicating similar bound state arrangements. **G)** Comparison of the LCDR3 contact site with the gp120 protein and N301-glycan between the I4 (upper) and DH270.3 (lower) structures. **H)** Comparison of the N332-glycan D1 arm antibody contacts between the I4 (upper) and DH270.2 (lower) structures.

A cluster of mutations, including an improbable V_H_ G110Y mutation, occur in and around the DH270.3 LCDR3, near the contact between a loop adjacent to the GDIK motif and the N301-glycan GlcNac base. Additional mutations, F93S, A94T, G95N, and S97A, result in remodeling of the N301-glycan interaction, shifting the position of the LCDR1 Y32 side chain for interaction with the first N301-GlcNac and interaction of N95 with the second N301-GlcNac (Figure 4G). In addition to the angular shift and LCDR3 related mutations, a G103S mutation in HCDR3 adds a hydrogen bond with the N332-glycan GlcNac-2 N-acetyl moiety (Supplemental Figure 8C). While DH270.3 mutations optimized interactions with the N301-glycan, the DH270.2 mutations focused instead on the D1 arm N332-glycan interaction. Of the three heavy chain mutations R98K, W101Y, and D115N, the D115N mutation showed the clearest shift in interaction. Loss of the negative charge at D115N allowed the asparagine side chain to shift away from the positively charged K98 to form an extensive hydrogen bonding network with the N332-glycan D1 arm (Figure 4H). This mirrored the R98T mutation in I3, where D115 was freed from its interaction with the positively charged V_H_ residue R98 to engage the N332-glycan. Together, analysis of the Env-bound DH270.2 and DH270.3 structures suggest each followed different paths of development with each targeting distinct sites. The greater breadth of the DH270.2 antibody compared to DH270.3 is likely a combined effect of improvement in the N332-glycan interaction in DH270.2 and the Y27S reversion in DH270.3.

### Maturation after I3: A complex series of mutations in I2 resolve epitope clashes

The next stage of development in the I3 clade of the DH270 clone led to the most substantial gains in Env binding affinity (Figure 1B) and HIV-1 neutralization breadth^21^. The I3 intermediate splits into the I2 intermediate and the mature DH270.1 antibody, both of which acquire mutations that are primarily focused on the LCDR3 region (Figure 5A and B). A notable feature in the cryo-EM reconstruction of the I2-Env complex is the well-resolved V1 loop with an altered configuration. Well-resolved density was also observed for the V1 loop N137-glycan, which contacted the I2 heavy chain S84R mutation site (Figure 5C and Supplemental Figure 9A). This mutation also occurred in the I4 intermediate which showed similar, albeit less clear, densities for a closed V1. Re-orientation of the V1 loop shifted R57 toward the GDIK motif, resulting in its interaction with D325 (Supplemental Figure 9B). The V1 loop closing over the V3 loop was similar to the UCA bound trimer structure with a kink in the loop introduced to accommodate R57 (Figure 5D). Quantification of the antibody orientation showed a rotation of the I2 V_H_/V_L_ away from the V1 loop to accommodate this loop rearrangement (Figure 5E).

**Figure 5.**
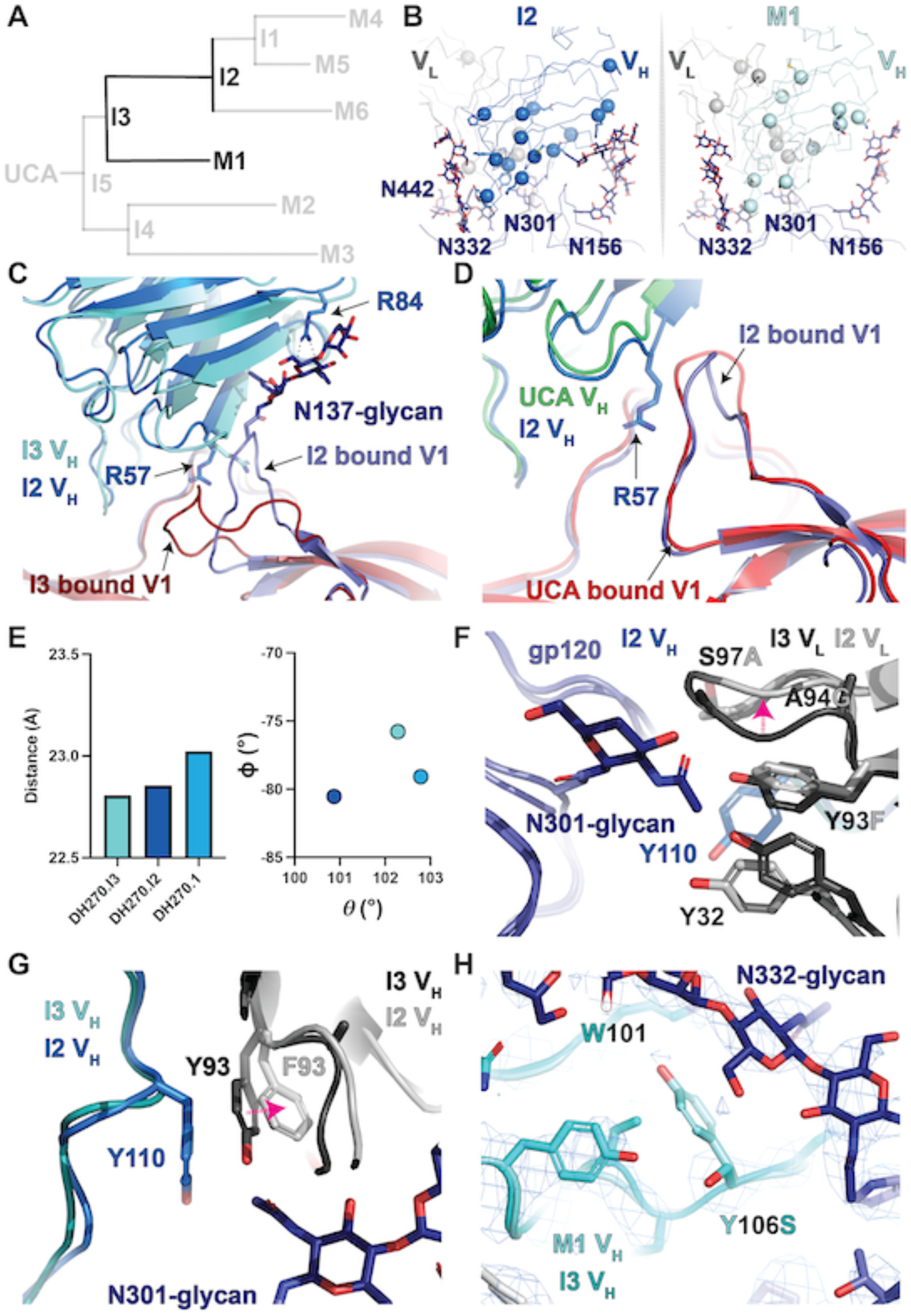
The I3 to I2 and DH270.1 path split. **A)** DH270 clonal tree highlighting the I3 and I2 intermediate antibodies and the mature DH270.1 (M1) antibody. **B)** Top views of the I2 and DH270.1 contact sites. Alpha-carbons (Cα) of heavy and light chain mutations are presented as spheres. **C)** Alignment (gp120 only) of I3 and I2 bound state structures highlighting the shift in gp120 V1 loop arrangement and contact between the I2 R84 side chain and the N137-glycan. **D)** Alignment (gp120 only) of UCA and I2 bound structures highlighting conformational similarity between the gp120 V1 loops. **E)** (right) Antibody Fab V_H_/V_L_ centroid distance from the gp120 epitope centroid. (left) Theta vs. phi plot for the I3, I2, and DH270.1 structures. The colors of the data points match the respective heavy chain colors in the structures. Side view comparison of the LCDR3 region epitope contact between I3 and I2 highlighting the shift in LCDR3 conformation (magenta arrow). **G)** Top view comparison of the LCDR3 region epitope contact between I3 and I2 highlighting the shift in the aromatic residue 93 rotamer (magenta arrow). **H)** Alignment (gp120 only) of I3 and DH270.1 bound state structures with DH270.1 map highlighting the Y106S mutation.

The I2 intermediate and DH270.1 both acquire several LCDR3 modifying mutations like those observed in DH270.3 (Figure 5F and G and Supplemental Figure 9C and D). In DH270.1 several mutations occurred at the LCDR3 base that were distal to the epitope, and together, they likely stabilized the LCDR3 region (Supplemental Figure 9E and F). Similar mutations occur in the I2 intermediate, most notably an F101C mutation spatially adjacent to the LCDR3 base that results in the formation of an additional disulfide bond in the antibody between residues 91 and 101 (Supplemental Figure 9D). This mutation is among the previously described minimal set of mutations needed to reach ∼90% of DH270.6 breadth^29^. A considerable number of mutations in and affecting LDCR3 were also a part of the minDH270 construct and have distinct effects on epitope interaction in the I2 intermediate bound structure. These included the V_L_ Y93F, A94G, and S97A that work in concert with V_H_ G110Y to cause a rearrangement in the LCDR3 region, creating a pocket in which the N301 GlcNac-1 N-acetyl moiety rests (Figure 5F). An important impact of the improbable HCDR3 G110Y mutation in creating this pocket is the shifting of the F93 rotamer via steric occlusion (Figure 5G). Additionally, an HCDR3 Y106S mutation occurs in DH270.1 that resulted in two effects. First, the loss of a hydrogen bond between the side chain of Y106 with the main chain of W101 and, second, release of the interaction of the Y106 side chain with the N332 glycan; together these may act to modulate the interaction of glycan N332 with HCDR3 by allowing it to engage the W101 side chain more effectively (Figure 5H). These results, together with the observed LCDR3 changes in DH270.3, suggest modifications of the interaction at this site were essential for affinity maturation.

### Maturation after I2: Maturation shifts to the epitope periphery

From the I2 intermediate the clone splits into two paths, both culminating in similar neutralization breadth, albeit with DH270.6 having the greatest potency (Figure 6A and B)^21^. The upper path from I2 leads to the I1 intermediate and then onto the mature DH270.4 and DH270.5 antibodies. Mutations in this I1 subclade were generally further from the antibody-antigen interactive surface with less clear structural impacts. In I1, two mutations with possible impacts on the LCDR2 loop, N62H and R65W, could affect N332-glycan D1 arm interactions (Figure 6C and Supplemental Figure 10A). This loop was, however, poorly resolved, limiting our ability to determine what, if any, impact these mutations have. In DH270.4, a V_H_ S85R mutation is positioned for interaction with the N156-glycan (Figure 6D and Supplemental Figure 10B). However, as was the case for the I1 intermediate mutations, this region is poorly resolved and the interaction cannot, therefore, be confirmed. The DH270.5 V_H_ and V_L_ acquired several mutations in proximity to the N301-glycan D arms (Figure 6E and Supplemental Figure 10C). This was also observed in the DH270.6 V_H_ and V_L_ (Figure 6F and Supplemental Figure 10D). In both cases the map densities left some question as to whether the N301-glycan D arm rests in this position. Additional mutations in DH270.6 occurred near the N442-glycan and the open state V1 loop (Figure 6G and H). Densities consistent with both the open and closed state V1 loop were apparent in the DH270.6 bound trimer (Figure 6H and Supplemental Figure 10E and F). Examination of these structures indicates mutations after I2 occurred primarily at regions more distal to the epitope, consistent with the ability of I2 to neutralize most of the viruses neutralized by DH270.6.

**Figure 6.**
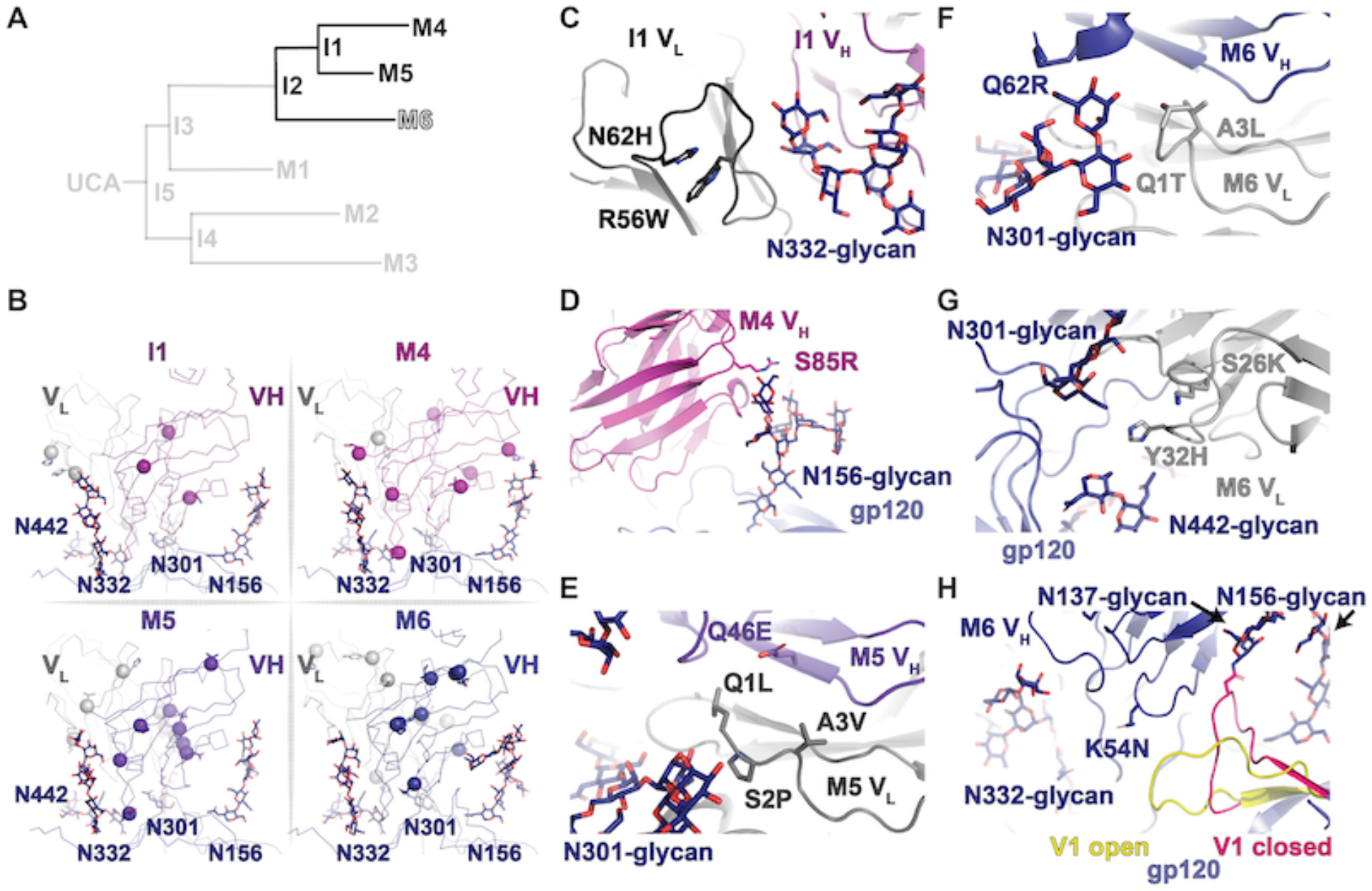
The I2 to I1, DH270.4, DH270.5, and DH270.6 transition. **A)** DH270 clonal tree highlighting the I2 and I1 intermediate antibodies and the mature DH270.4 (M4), DH270.5 (M5), and DH270.6 (M6) antibodies. **B)** Top views of the I1, DH270.4, DH270.5, and DH270.6 contact sites. Alpha-carbons (Cα) of heavy and light chain mutations are presented as spheres. **C)** Structure of the I1 V_H_/V_L_ contacts with the N332-glycan D1 arm showing the positions of the V_L_ N62H and R56W mutations. **D)** Structure of the DH270.4 heavy chain showing the position of possible contact between the S85F mutation and the N156-glycan. **E)** Structure of the DH270.5 V_H_/V_L_ mutations proximal to the N301-glycan. **F)** Structure of the DH270.6 V_H_/V_L_ mutations proximal to the N301-glycan. **G)** Structure of the DH270.6 V_L_ mutations proximal to the N442-glycan. **H)** Structure of the DH270.6 V_H_ showing two different states of the gp120 V1 loop.

Together, structural determination of the entire DH270 clone revealed an intricate process through which HIV-1 neutralization breadth was developed in the V3-glycan supersite directed DH270 clone. The early improbable G57R and S27Y mutations of the I5 intermediate had profound impacts on antibody orientation that are likely essential for how the antibodies developed downstream. Development of two distinct clades from I5 to the I3 and I4 intermediates resulted in both acquiring improvements in N332-glycan interaction further highlighting the importance of N332-glycan engagement for the DH270 clone. Each, however, targeted distinct sites of the glycan, with I3 stabilizing the distal D1 arm and I4 stabilizing the GlcNac base. Downstream maturation from I4 to DH270.2 suggests this was not an ideal solution to acquiring N332-glycan interaction, as DH270.2 acquires mutations like those in I3 for the N332-glycan D1 arm. Post-I3 development, leading to I2 and DH270.1 along with DH270.3, all show a considerable number of mutations in the LDCR3 and adjacent residues. While DH270.3 acquires several mutations critical for neutralization, including G110Y and S94A, the selection of the Y27S reversion may have limited its breadth. Where DH270.1 acquired stabilizing LCDR3 mutations, I2, which developed considerable neutralization breadth, modified the LCDR3 conformation, stabilizing a N301-glycan base contact point. This was, critically, mediated by at least five residues, each of which alone may have provided limited affinity gain. In the case of DH270.3, had it first stabilized the N332-glycan D1 arm, the limited gains associated with optimizing LCDR3 site might have been tractable. Paratope modifications downstream from the I2 intermediate were distant from the primary epitope and likely acted to fine tune affinity, breadth, and potency. The minDH270 antibody^29^ required only mutations from I5 to I2, consistent with this observation. These results show that selection during affinity maturation can be narrowed to a series of structural bottlenecks with the developing antibody clone demonstrating optimal and suboptimal solutions as the clone splits into distinct paths.

### Envelope structural evolution optimizes DH270 clone affinity maturation

The evolution of HIV-1 neutralization breadth in the DH270 clone occurred in the backdrop of co-evolution with the virus, with key changes in the Env protein driving antibody maturation, and vice versa^21^. We first examined how the DH270 epitope evolved during infection in the CH848 HIV-1 infected individual (Supplemental Figure 11). The first DH270 clone antibodies appeared between 793- and 1304-days post infection coinciding with a reduction in Env V1 loop length by 18-19 amino acids and a G300N substitution in Env at a location proximal to the DH270 epitope that appeared at day 699 post infection (Figure 7A). The early I5 and I3 intermediates recognize only the Envs that harbor these shorter V1 loops and the G300N mutation. Consistent with Env selection driven by the earlier DH270 clone antibodies I5 and I3 (Supplemental Figure 12A), autologous viruses with longer V1 loops and G300 re-emerge 1304-1431 days post infection. Ability of the DH270 clone to neutralize viruses with both longer V1 loop lengths and G300 appears at the I2 intermediate (Figure 7A, Supplemental Figure 12). Recognition of longer V1 loops and residue 300 mutations in CH848 quasispecies is also mirrored in DH270 antibody recognition of heterologous viruses with these features, contributing to the enhanced breadth of I2 and DH270.6 as compared to I5 and I3^21^.

**Figure 7.**
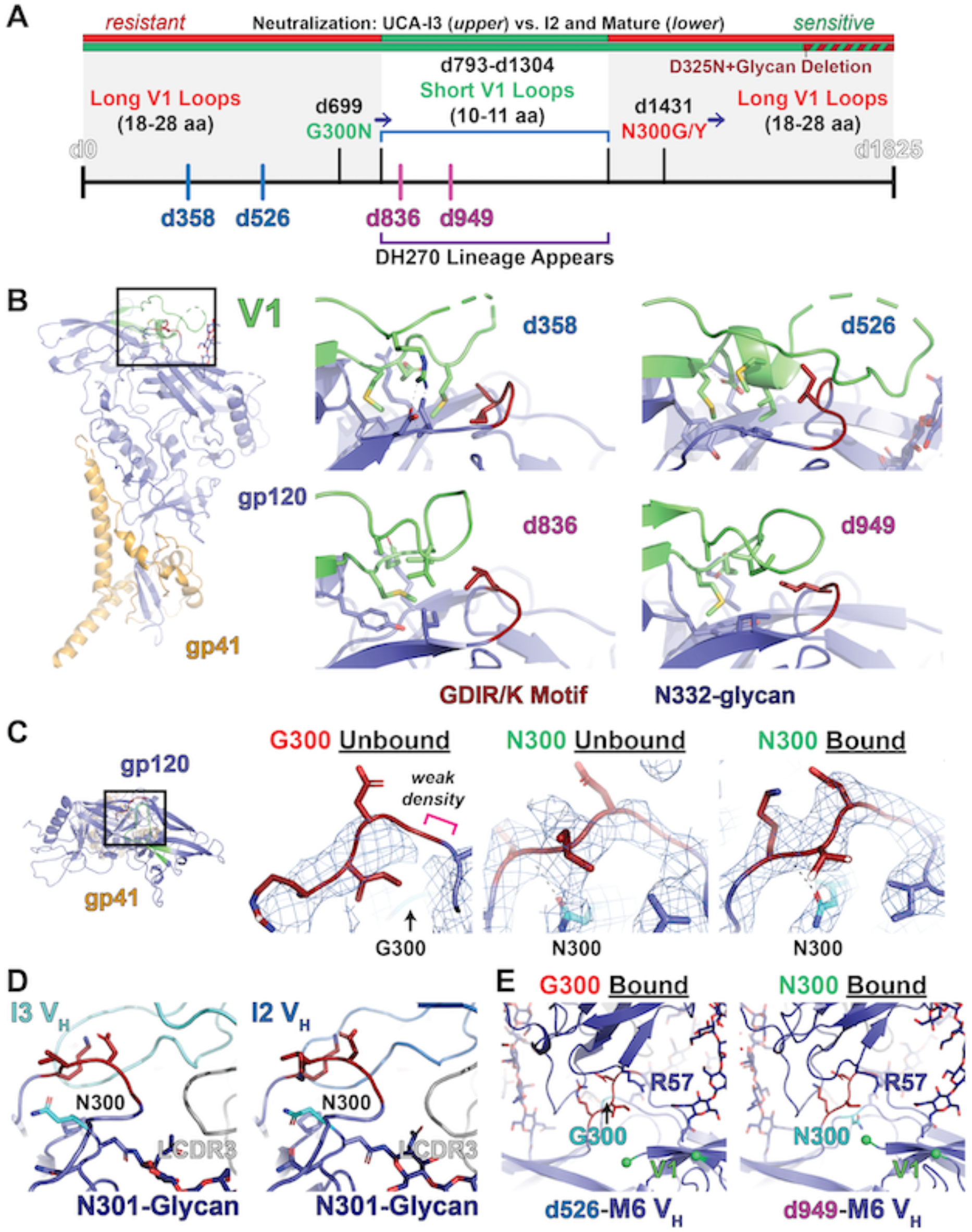
Envelope-DH270 clone structural coevolution. **A)** Timeline of antibody and Env coevolution. The upper boxes indicate whether Envs at each time point are sensitive (green) or resistant to early intermediates (upper) and later intermediates/mature (lower) clonal members. Periods of infection at which long V1 loops are dominant are shown in grey with key mutation time points are highlighted along the timeline axis. Env structures were determined for sequences isolated at days 358, 526, 836, and 949 post infection highlighted on the timeline axis with period at which the D270 clone appeared highlighted. **B)** (left) Side view of a single gp120/gp41 protomer highlighting the V1 and GDIR/K motif. (right) V1 loop structures of each Env isolate. **C)** (left) Top view of a single gp120/gp41 protomer highlighting the V1 and GDIR/K motif. Maps and structure fits for the unliganded d358 (middle left), unliganded d949 (middle right), and I3 bound d949 (right) structures. **D)** The I3 (left) and I2 (right) bound structures showing similarity in epitope conformation. **E)** The DH270.6 bound d526 structure (left) and DH270.6 bound d949 structure (right) showing similarity in epitope conformation.

Based on these two early intermediate resistance features, we selected Envs from days 358 and 526, having both G300 and long hypervariable V1 loops of 18 and 28 amino acids, respectively, and Envs from days 836 and 949, having both N300 and short hypervariable V1 loops of 11 amino acids, to identify structural hurdles to DH270 antibody interaction. We first examined binding of the clone antibodies to each Env SOSIP ectodomain by ELISA (Supplemental Figure 13). Consistent with previous observations, the UCA displayed only binding to the d949 construct with I5 binding the d949 and the d836 constructs. While I3 displayed binding to the d358 construct, I4 did not. Consistent with differences in the early maturation of these clones, the mature DH270.2 and DH270.3 antibodies bound weaker to the d358 construct compared to I2, I1, and DH270.4-6. Neither DH270.2 or DH270.3 bound the d526 construct while measurable binding to I2, I1 and DH270.4-6 was observed, though the interaction was weaker than to the other three constructs. Despite poor sequence similarity, examination of the V1 loop regions of each unliganded Env structure demonstrated a structurally conserved hydrophobic core and variable hydrogen bonding and/or salt bridge formation that ensures coupling of the V1 loop to the GDIR/K motif. The d358 Env V1 displayed considerable disorder in a portion of the loop with flanking hydrophobic and salt bridge interactions ensuring the loop remained closed over the GDIR/K motif (Figure 7B and Supplemental Figure 14A). The d526 Env contained a similar hydrophobic core and displayed more order despite its longer length. This was due to direct V1 loop interaction with the nearby N332-glycan (Figure 7B and Supplemental Figure 14B). The d836 and d949 Env V1 were markedly shorter but still retain the hydrophobic core and closed state position of the earlier viruses (Figure 7B and Supplemental Figure 15). The d836 loop displayed a kink in the loop near the GDIR/K motif leading to somewhat greater exposure of the motif (Figure 7B and Supplemental Figure 15C). Together, these results indicate a shared role of the V1 for protecting the neutralization sensitive GDIR/K motif that is mediated by a conserved hydrophobic core.

We next examined the effect on the DH270 epitope of the Env G300N substitution that is associated with clone initiation. The GDIR/K motif is proximal to position 300, forming a paired, broken beta sheet secondary structure. Visual inspection of the GDIR/K motif in the G300-containing d358 and d526 Env maps suggested greater disorder in the broken sheet structure compared to that of the N300-containing d836 and d949 Env maps (Figure 7C). N300 formed a hydrogen bonding contact with the I326 backbone of the GDIK motif in the unbound state. This interaction is retained in the bound state structures for the early intermediates with little change in the GDIK motif conformation (Figure 7C). These results suggest the G300N stabilized the DH270 epitope enabling the UCA and early intermediates to bind and neutralize the virus. Mutations at residue N300 begin to occur at 1431 days post infection (Supplemental Figure 11) at which point glycine and tyrosine variants co-circulate with asparagine variants. This occurs around the time that the LCDR3 mutations proximal to Env residue 300 occur in I2, DH270.1, and DH270.3, suggesting the evolving virus selected for these mutations in the clone. Comparison of the Env-bound I3 and I2 structures indicated little overall change in their epitope configuration (Figure 7D). Additionally, structures of DH270.6 bound to d526 Env containing G300 and d949 Env containing N300 show few changes in the overall epitope conformation (Figure 7E and Supplemental Figure 14B). As the principal effect of the LCDR3 mutations was rigidification of the LCDR3 and closer contact with the N301-glycan base, with little change to the bound state configuration in the unliganded or bound states, these mutants likely enhanced the interaction by compensating for increased epitope flexibility. Additional coevolving Env sites outside the V1 loop and position 300 included positions 325 to 327 in the ^324^GDIR/K^327^ motif and position 328 (Supplemental Figure 11 right). Mutations also occurred that eliminated the N301, N332, and N442 glycan sequons beginning at 1304 days post infection. Autologous viruses that were resistant to the mature DH270.4 were predominantly enriched for the D325N substitution, but also had loss of glycans at 301, 332 and/or 442. A clear structural impact for D325N was not observed while the close contacts in all structures with the glycans suggest their loss would indeed have major impacts on DH270 antibody binding. Together, these results show that virus evolution initiated DH270 clonal evolution by stabilizing the GDIR/K epitope through G300N and removing the long V1 loop protection of the conserved V3 loop GDIR/K motif. The return of these protective features then spurred maturation of the clone toward stabilization of the paratope to combat enhanced flexibility, resulting in greater neutralization breadth.

## Discussion

Here we have studied a series of cryo-EM structures and correlated them with antibody affinity outcome in a prototype V3-glycan HIV-1 broadly neutralizing antibody B cell clone. We demonstrated the early G57R and S27Y mutations required for acquisition of heterologous breadth by the I5 intermediate antibody had profound impacts on antibody orientation that likely affected downstream antibody development. Clonal differentiation into the more mutated I3 and I4 intermediates resulted in both acquiring improvements in N332-glycan interaction but targeting different arms of the branched glycan. Downstream maturation from I4 to DH270.2 suggested this was not an ideal solution to optimizing N332-glycan interaction, as DH270.2 acquires mutations like those in I3 for the N332-glycan D1 arm. If DH270.2 shows that I4 selection was not ideal, DH270.3 shows the trouble with moving on from a critical juncture too soon. Post-I3, development to I2 and DH270.1 along with DH270.3 all showed a considerable number of mutations in the LDCR3 and adjacent residues with I2 acquiring the greatest neutralization breadth and potency. Paratope modification downstream from the I2 intermediate appeared further from the primary epitope and likely acted to fine tune affinity, breadth, and potency. These results showed staged, site-specific development determined the fate of downstream DH270 clonal affinity maturation.

A recent study of a broadly neutralizing influenza hemagglutinin targeting antibody B cell clone suggested affinity maturation involves developing a diverse pool of a potential solutions from which further optimization could induce a broad response^4^. Consistent with that study, here we found that differing mutations solved similar problems presented by the pathogen. However, these solutions were not necessarily equally effective, and the developing clone appeared to evolve toward optimal solutions to gain affinity as evinced by clade mutation similarity. What supports this antibody “pluripotency” hypothesis developed based on an influenza bnAb^4^ creates a vexing problem for vaccine design. As specific endpoints in clones are desired for their breadth and potency properties, careful steering of developing responses towards these endpoints and away from less productive solutions for neutralization breadth will likely be required.

Our results indicated that solutions to epitope-paratope pairing issues, rather than occurring together, resulted in clade diversification. Seen from the perspective of the entire DH270 clone, failure to acquire an optimal solution limits further affinity maturation as was the case for clonal members DH270.1 and DH270.3. Thus, failing to acquire recognition solutions to specific problem sites early in bnAb lineage maturation is likely to hinder favorable development of later lineage member neutralization breadth. In the context of the DH270 clone, vaccination that fails to induce selection of the I5 G57R and S27Y mutations may limit acquisition of the I3 R98T and L48Y mutations. Further, our results demonstrated how off-track mutations can lead clone development to suboptimal breadth. Knowing which off-target responses are likely to occur and which portions of the epitope may be involved in selecting for mutations that lead the clone off-track allows us to select immunogen modifications with greater precision, ensuring modifications produce a positive differential affinity gain favoring the desired mutation(s). Unlike I3, which contained two critical mutations likely to result in affinity gain individually, I2 acquired multiple LCDR3 mutations that worked together to modify interaction with the Env. The need to induce multiple mutations that may do little to improve affinity individually will likely require thinking beyond immunogen properties to issues related to adjuvant, vaccination timing, and the number and identity of boosting immunogens.

Beyond providing a time-resolved, atomic-level definition of the development of the HIV-1 bnAb DH270 clone over several years, the insights from this study provides a clear roadmap to precision, clone-based immunogen development. By isolating distinct structural elements at the antibody-antigen interface, sites of gain can be established, and immunogens can be developed to sequentially solve structural problems associated with each site individually. By shifting focus to specific, manageable steps, rational affinity design can proceed through a series of well-defined targets rather than attempting to implement a speculative set of final immunogen designs lacking clear data.

## Methods

### Recombinant antibody production

Expi293 cells (ThermoFisher Cat No. A14527) were diluted to a final volume of 0.5L at a concentration of 2.5×10^6^ cells mL^−1^ in Expi293 media. Four hundred micrograms of heavy chain and light chain plasmid were complexed with Expifectamine (ThermoFisher Cat No. A14526) and added to the Expi293 cells. On day five, cells were cleared from transfection cell culture media by centrifugation and 0.8 μM filtration. The supernatant containing recombinant antibody was incubated with protein A resin (ThermoFisher) overnight at 4 °C. The protein A resin was collected by centrifugation and culture supernatant was removed. The resin was washed with 25 mL of phosphate-buffered saline (PBS) containing a total of 340 mM NaCl. Thirty mL of 10 mM glycine pH 2.4, 150 mM NaCl were used to elute the antibody off of the protein A resin. The pH of the eluted antibody solution was increased to approximately 7 by the addition of 1M Tris pH 8.0. The antibody solution was buffer exchanged into PBS with successive rounds of centrifugation, filtered, and stored at −80 °C.

### Recombinant HIV-1 envelope SOSIP gp140 production

CH848 SOSIP gp140 envelope production was performed with Freestyle293 cells (ThermoFisher Cat No. R79007). On the day of transfection, Freestyle293 were diluted to 1.25×10^6^ cells/mL with fresh Freestyle293 media up to 1 L total volume. The cells were co-transfected with 650 μg of SOSIP expressing plasmid DNA and 150 μg of furin expressing plasmid DNA complexed with 293Fectin (ThermoFisher Cat No. 12347019). On day 6 cell culture supernatants were harvested by centrifugation of the cell culture for 30 min at 3500 rpm. The cell-free supernatant was filtered through a 0.8 μm filter and concentrated to less than 100 mL with a single-use tangential flow filtration cassette and 0.8 μm filtered again. Trimeric Env protein was purified with monoclonal antibody PGT145 affinity chromatography. PGT145-coupled resin was packed into Tricorn column (GE Healthcare) and stored in PBS supplemented with 0.05% sodium azide. Cell-free supernatant was applied to the column at 2 mL/min using an AKTA Pure (GE Healthcare), washed, and protein was eluted off of the column with 3M MgCl_2_. The eluate was immediately diluted in 10 mM Tris pH 8, 0.2 μm filtered, and concentrated down to 2 mL for size exclusion chromatography. Size exclusion chromatography was performed with a Superose 6 16/600 column (GE Healthcare) in 10 mM Tris pH 8, 500 mM NaCl. Fractions containing trimeric HIV-1 Env protein were pooled together, sterile-filtered, snap frozen, and stored at −80 °C.

### Surface Plasmon Resonance

The SPR binding curves of the DH270 lineages Fabs against CH848 DS SOSIP Trimer were obtained using either a Biacore S200 instrument (Cytiva) in HBS-N 1X running buffer or a Biacore 3000 instrument (Cytiva) in PBS 1X pH7.4. For DH270 Fabs I5.6, I3.6, I2.6, and DH270.6 affinity measurements, biotinylated CH848 DS SOSIP Trimer was immobilized onto a CM3 sensor chip via streptavidin to a level of 280-290RU. Using the single cycle injection type, six sequential injections of the Fabs diluted from 50 to 1500nM were injected over the immobilized SOSIP Trimer at 50uL/min for 120s per concentration followed by a dissociation period of 600s. The Fabs were regenerated with a 20s pulse of 25mM NaOH at 50uL/min. Results were analyzed using the Biacore S200 Evaluation Software (Cytiva). A blank streptavidin surface as well as buffer binding were used for double reference subtraction to account for non-specific antibody binding and signal drift. Curve fitting analyses were performed using the 1:1 Langmuir model with a local Rmax. For DH270UCA3 Fab, biotinylated CH848 DS SOSIP Trimer was immobilized onto a CM5 sensor via streptavidin to a level of 290-300RU. DH270UCA3 Fab was diluted from 500 to 3000nM and each concentration was injected over the SOSIP Trimer surface for 300s using the K-inject injection type at 50uL/min. A 600s dissociation period was followed by surface regeneration with a 20s pulse of 50mM NaOH. Results were analyzed using the BIAevaluation software (Cytiva). A blank streptavidin surface and buffer binding were used for double reference subtraction. Curve fitting analysis of DH270UCA3 Fab was performed using the 1:1 Langmuir model with a local Rmax. The reported binding curves for all Fabs are representative of one data set.

### Cryo-EM Sample Preparation

The CH848 SOSIP trimer complexes were prepared using a stock solution of 2 mg/ml trimer incubated with a six-fold molar excess of Fab. To prevent interaction of the trimer complexes with the air-water interface during vitrification, the samples were incubated in 0.085 mM *n*-dodecyl β-D-maltoside (DDM). Samples were applied to plasma-cleaned QUANTIFOIL holey carbon grids (EMS, R1.2/1.3 Cu 300 mesh) followed by a 30 second adsorption period and blotting with filter paper. The grid was then plunge frozen in liquid ethane using an EM GP2 plunge freezer (Leica, 90-95% relative humidity).

### Cryo-EM Data Collection

Cryo-EM imaging was performed on a FEI Titan Krios microscope (Thermo Fisher Scientific) operated at 300 kV. Data collection images were acquired with a Falcon 3EC or K3 Direct Electron Detector operated in counting mode with a calibrated physical pixel size of 1.08 Å with a defocus range between −1.0 and −3.5 µm using the EPU software (Thermo Fisher Scientific). No energy filter or C_s_ corrector was installed on the microscope. The dose rate used was ∼0.8 e^−^/Å^2^·s to ensure operation in the linear range of the detector. The total exposure time was 60 s, and intermediate frames were recorded every 2 s giving an accumulated dose of ∼42 e^−^/Å^2^ and a total of 30 frames per image.

### Data Processing

Cryo-EM image quality was monitored on-the-fly during data collection using automated processing routines. For datasets DH270.I3, DH270.I2, DH270.6, DH270.I4 and DH270.2, data processing was performed within cryoSPARC^26^ including particle picking, multiple rounds of 2D classification, *ab initio* reconstruction, homogeneous map refinement and non-uniform map refinement. The rest of the datasets were further processed outside of cryoSPARC to improve map quality as described next. Movie frame alignment was carried out using UNBLUR^27^, and CTF determination using CTFFIND4^28^. Particles were picked automatically using a Gaussian disk of 80 Å in radius as the search template. All the picked particles were extracted with box size of 384 pixels and down scaled to 192 pixels (binning 2) and subjected to 8 rounds of refinement in cisTEM^29^, using an *ab-initio* model generated with cryoSPARC^26^. The distribution of scores assigned to each particle by cisTEM showed a clear bi-modal distribution and only particles in the group containing the higher scores were selected for further processing. The clean subset of particles were re-extracted without binning and subjected to 10 iterations of local refinement followed by 5 additional iterations using a shape mask generated with EMAN2^30^. Per-particle CTF refinement was then conducted until no further improvement in the FSC curve was observed. At this point, particle frames were re-extracted from the raw data and subjected to per-particle motion correction using all movie frames and applying a data-driven dose weighting scheme^31^. The new locally aligned particles were exported to cryoSPARC to conduct an additional round of homogeneous refinement and non-uniform refinement.

For elbow angle analysis with DH270.I3, I4, and DH270.5, local focus refinement with the clean particles after symmetry expansion was done in cryoSPARC. Then the particles together with corresponding refinement parameters were then exported into RELION 3.1 to perform 3D classification without alignment using T=25.

### Cryo-EM Structure Fitting

Structures of each antibody were prepared using Modeller^32^ and available Fab crystal structures^21,30^. The SOSIP Env structures were prepared using modeler and the DH270.6 bound CH848 day 949 trimer (PDB ID 6UM6 chains A and B)^22^. Coordinates for VRC01 from PDB ID 3NGB^33^ (chains H and L). Structure fitting of the cryo-EM maps was performed in ChimeraX^34^ using the Isolde application^35^. Structure fit and map quality statistics were determined using MolProbity^36^ and EMRinger^37^, respectively. Glycans were fit using a combination of the sharpened and gaussian filtered maps (1.0-1.5 standard deviation). Structure and map analyses were performed using a combination of PyMol^38^ and ChimeraX with theta and phi angles calculated using VMD^39^.

#### ELISA

Antibody binding to SOSIP proteins was tested using 384-well ELISA plates were coated with 2mcg/ml of antibody in 0.1 M sodium bicarbonate overnight at 4°C. Plates were washed with PBS/0.05% Tween-20 and blocked with assay diluent (PBS containing 4% (w/v) whey protein/15% Normal Goat Serum/0.5% Tween-20/0.05% Sodium Azide) at room temperature for 1 hour. Purified SOSIP protein was added in 2-fold serial dilutions starting at 10mcg/ml for 1 hour followed by washing. Binding of SOSIP to the coated antibodies was detected using biotinylated human antibody PGT151 at 0.125mcg/ml for 1 hour. PGT151 was washed and followed by streptavidin-HRP (Thermo Scientific # 21130) 1:30,000 for 1 hour. Plates were washed 4 times and developed with TMB substrate (SureBlue Reserve-#5120-0083) followed by 1M HCl to stop the reaction. Finally the absorbance at 450 nm (OD_450_) was determined with a 384 well plate reader (Molecular Devices, Spectramax 384 plus).

### CH848 longitudinal Env analyses

Env sequences used for signature analysis are from Bonsignori et al^21^. The webtool AnalyzeAlign at the Los Alamos HIV Database was used for generating sequence logos. Autologous neutralization data are from Bonsignori et al.^21^ measured on 90 autologous Env pseudoviruses including the TF and representative longitudinal Envs sampled between 78 to 1720 days post infection. While intermediates in the previous study were inferred with 5 clone members instead of 6 as in this current study, intermediates I5, I3 and I2 correspond very closely to IA4, IA2 and IA1, respectively in Bonsignori et al.^21^

## Figures

**Supplemental Figure 1.**
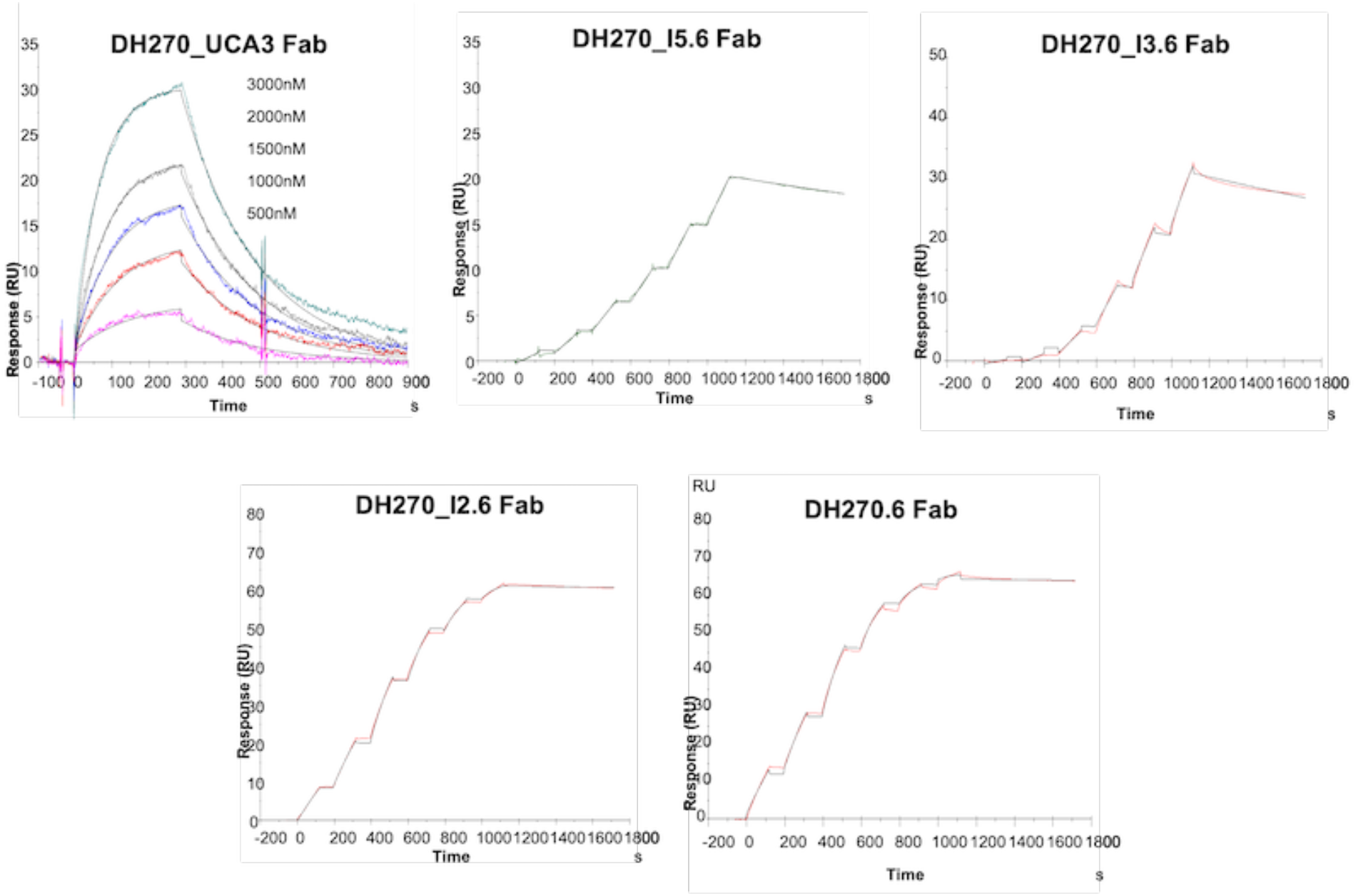
Surface plasmon resonance (SPR). Binding curves are overlaid with curves fits used to derive affinities for antibody Fabs leading from the UCA to the mature DH270.6 listed in Figure 1.

**Supplemental Figure 2.**
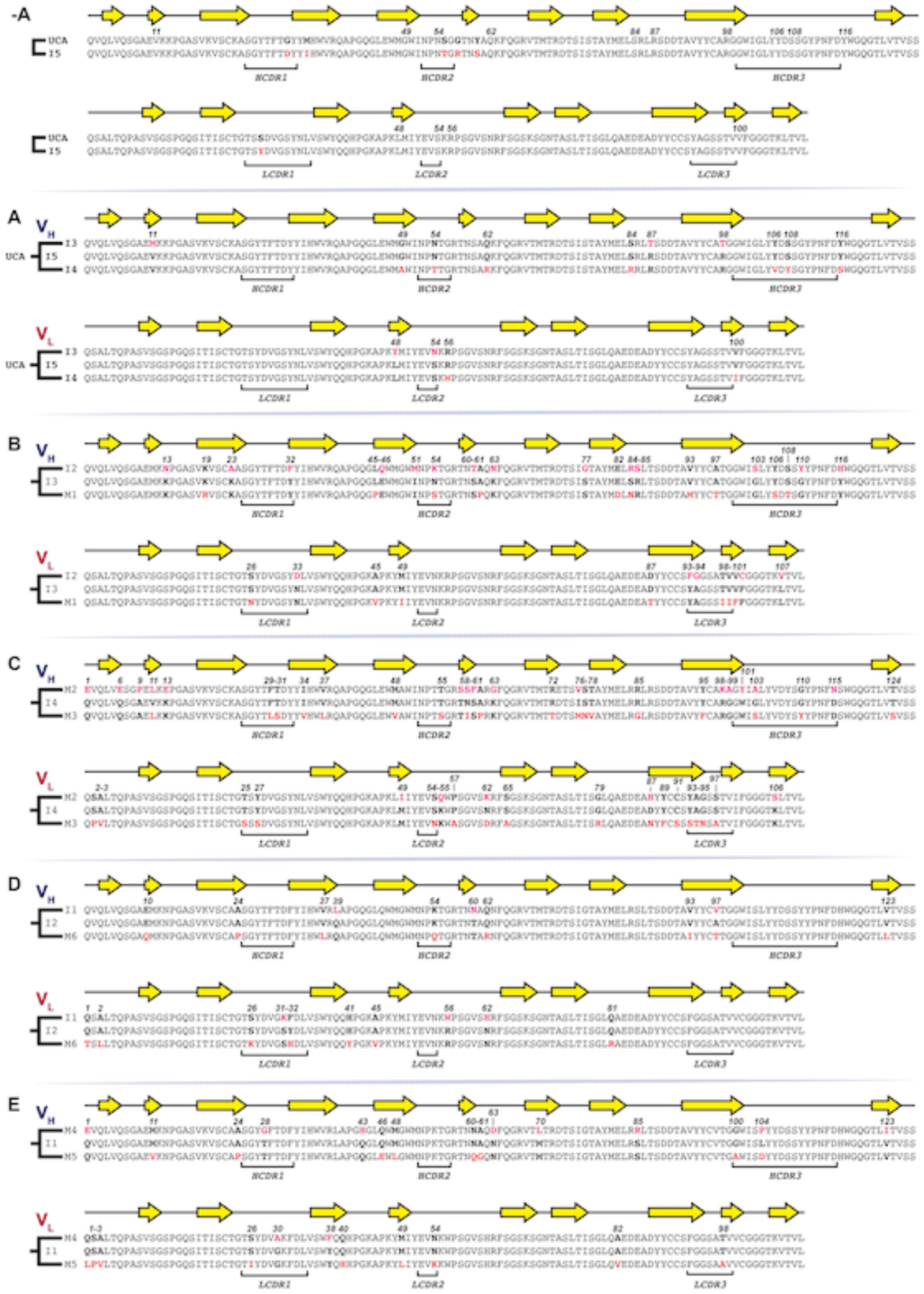
Sequence alignments for the DH270 clonal clone. Sequence alignments at each path split with secondary structure and mutations highlighted in bold.

**Supplemental Figure 3.**
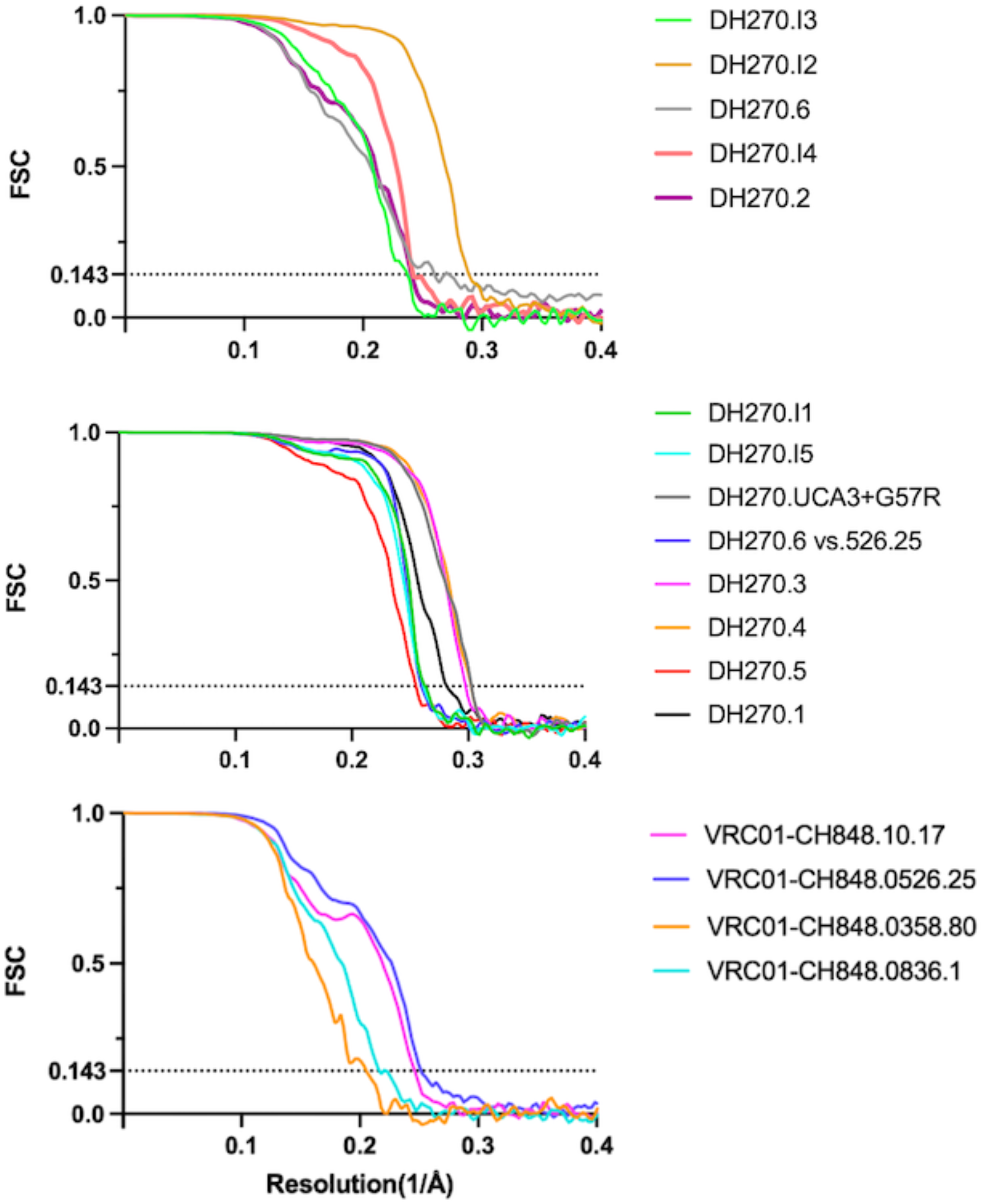
Fourier shell correlation (FSC) plots for DH270 clonal clone antibody Fab bound maps, set one. Gold-standard FSC curves calculated from two independently refined half-maps. The dotted line indicates FSC = 0.143.

**Supplemental Figure 4.**
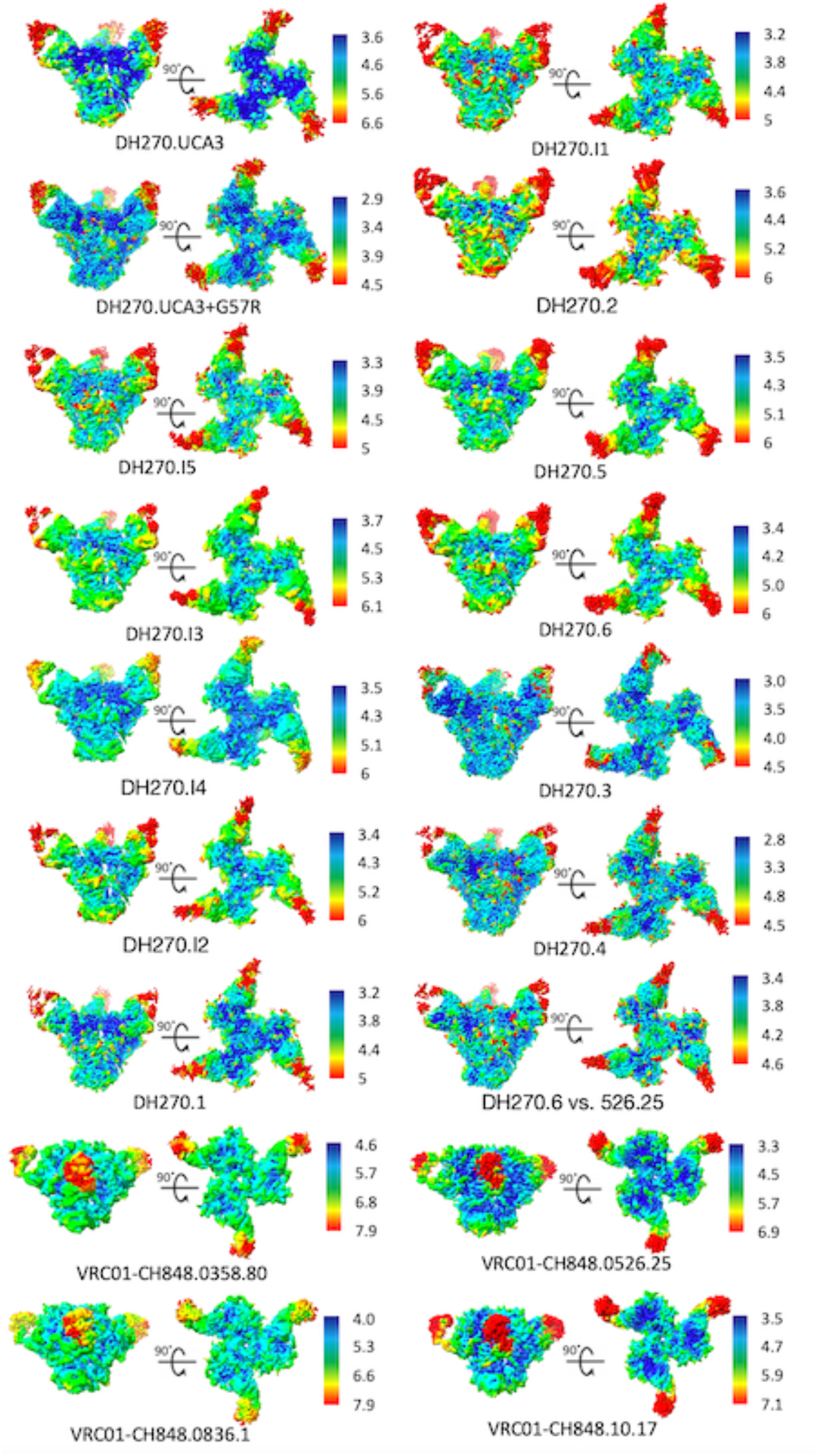
Fourier shell correlation (FSC) plots for DH270 clonal clone antibody Fab bound maps, set two. Gold-standard FSC curves calculated from two independently refined half-maps. The dotted line indicates FSC = 0.143.

**Supplemental Figure 5.**
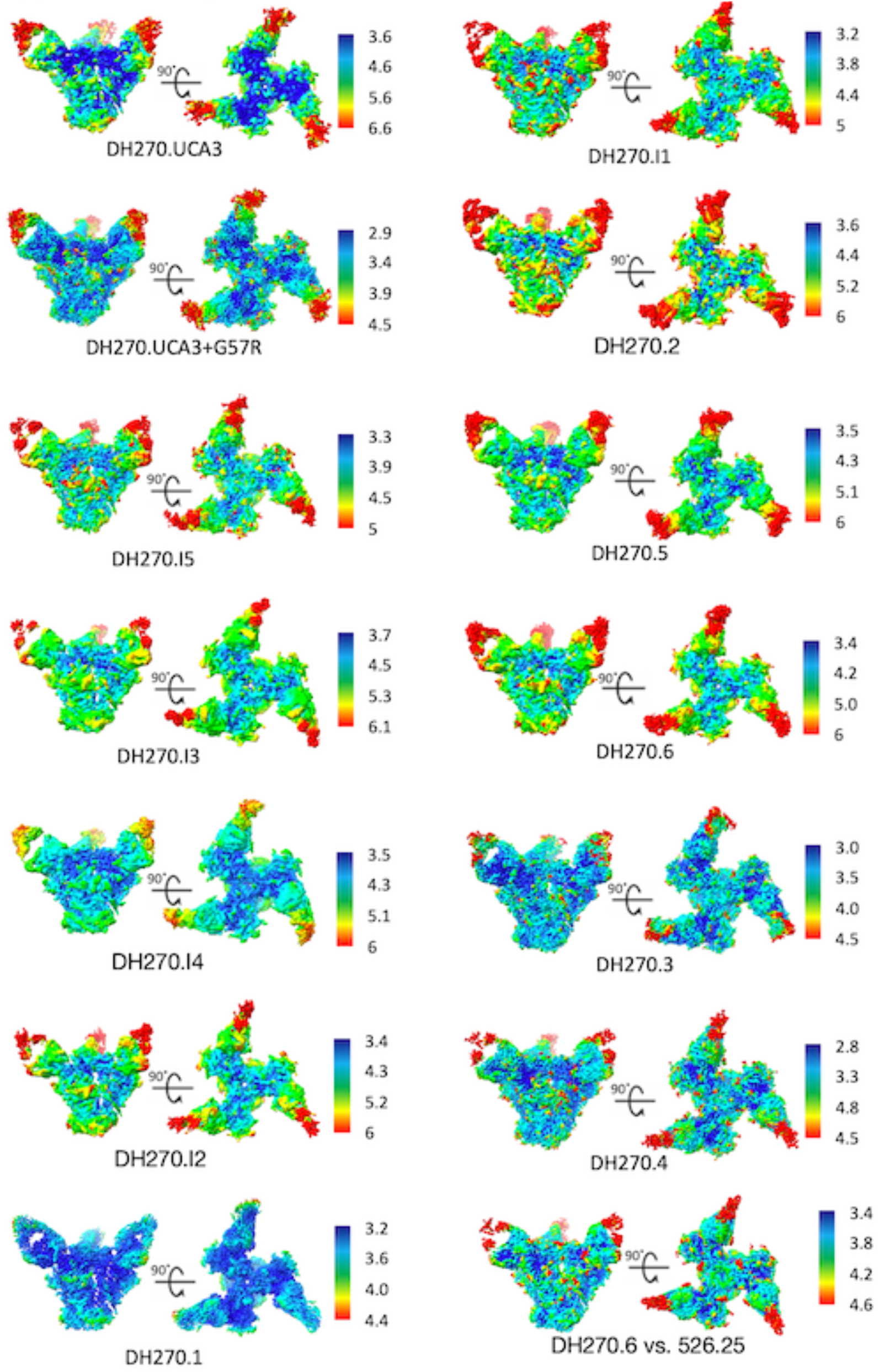
Local resolution for the DH270 clonal clone antibody Fab bound structures, set two. Refined cryo-EM maps for each complex colored by local resolution.

**Supplemental Figure 6.**
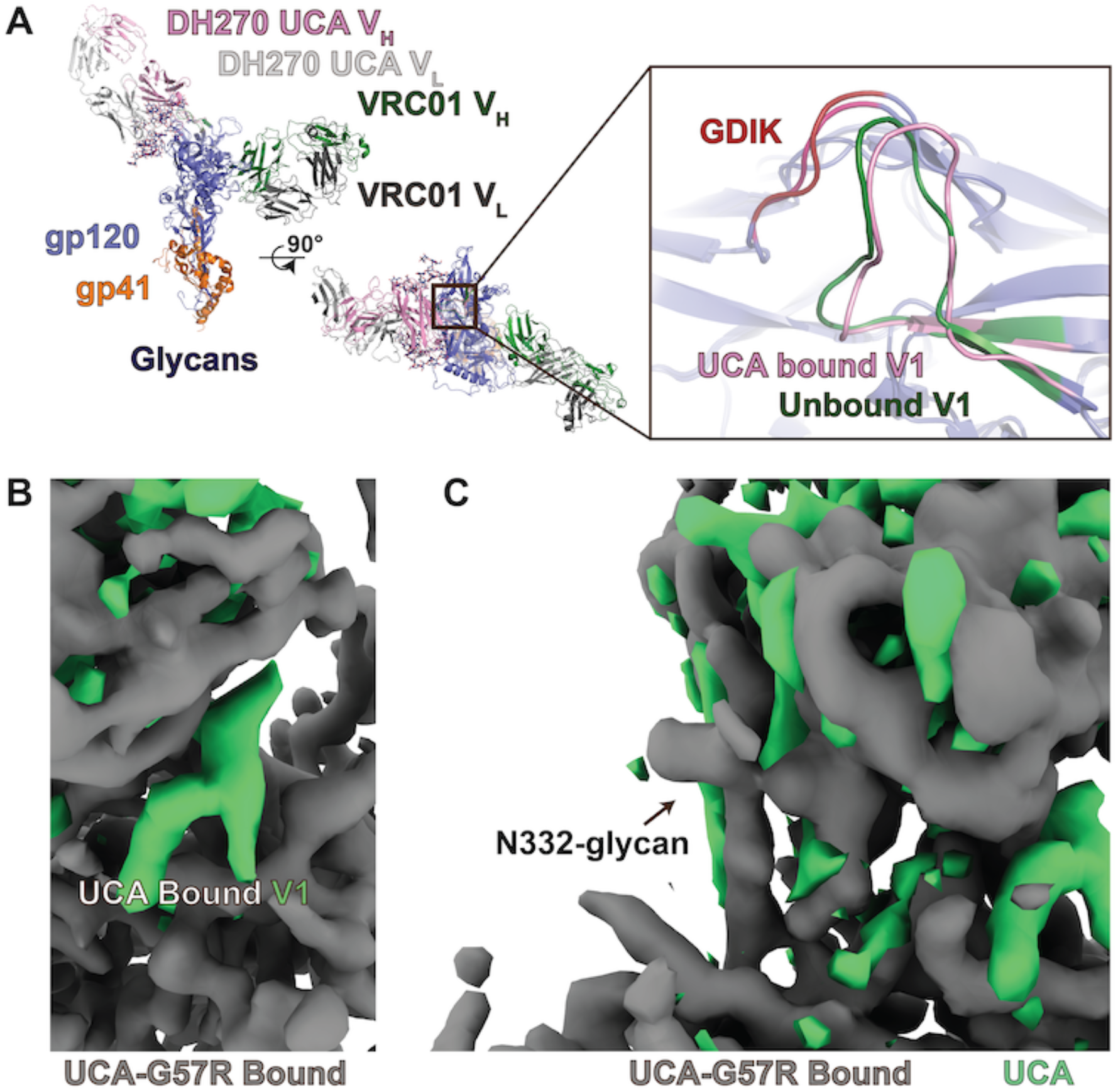
Changes in V1 loop configuration induced by heavy chain G57R mutation early in DH270 clonal development (UCA to I5). **A)** Aligned gp120/gp41 domains of the DH270 UCA and VRC01 bound CH848 d949 structures highlighting similarity between the liganded (UCA bound structure) and unliganded (VRC01 bound structure) epitope V1 loop conformations. **B)** The UCA+G57R map (grey) overlaid with a UCA difference map (green) highlighting V1 loop differences. **C)** The UCA+G57R map (grey) overlaid with a UCA difference map (green) highlighting the shift in antibody position.

**Supplemental Figure 7.**
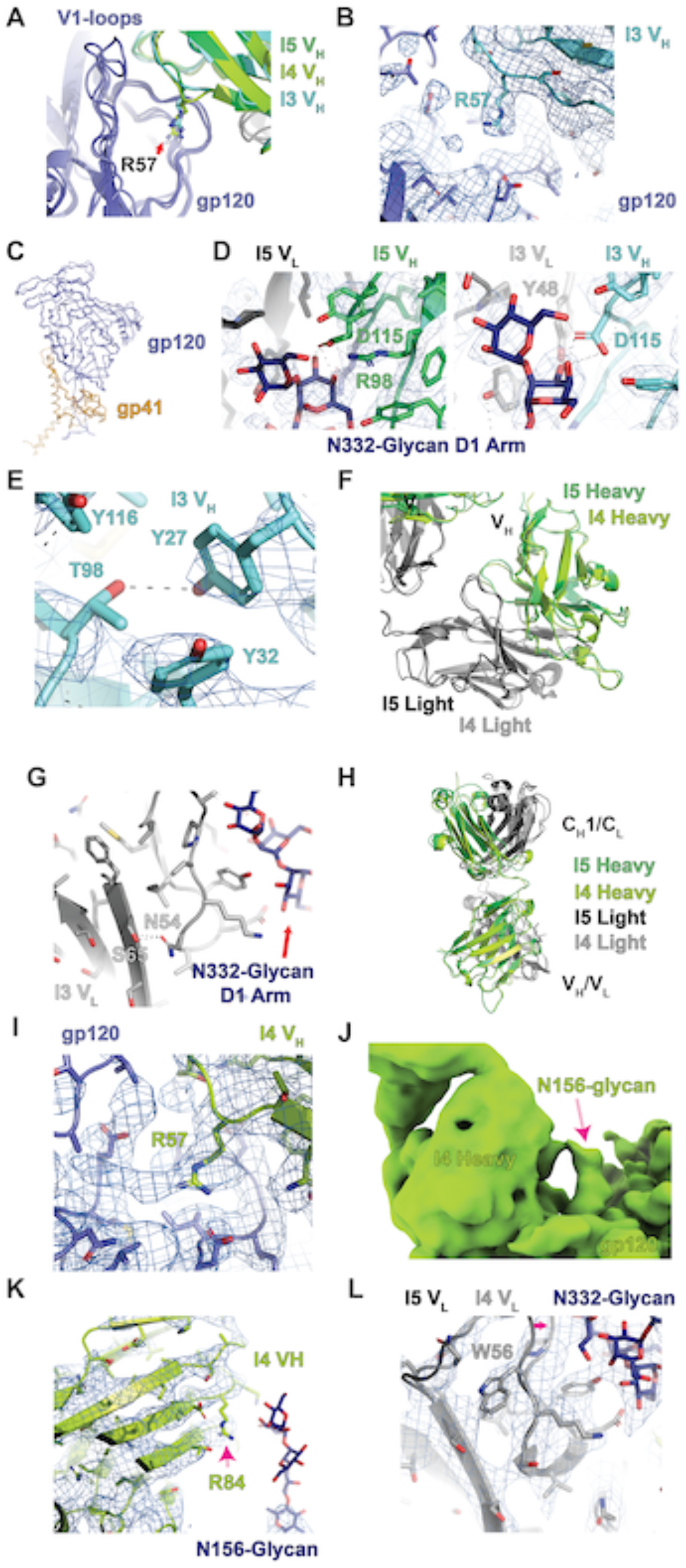
The I5 path split structures: I3 and I4. **A)** gp120 V1 loops displaced by residue R57 of I5, I4, and I3. **B)** Map and fit for the I3 intermediate showing the regions around residue R57. **C)** Alignment of the I5, I4, and I3 bound gp120/gp41 domains. **D)** (left) Map and fit for the I5 intermediate of the regions around residues R98 and D115 showing interactions with the N332-glycan D1 arm. (right) Map and fit for the I3 intermediate of the regions around the R98T and D115 positions showing improved interactions with the N332-glycan D1 arm. **E)** Map and fit of the I3 Fab V_H_ showing interaction between Y27 and T98. **F)** Comparison of the I5 and I4 intermediate antibody bound state from alignment of each structure’s gp120 domain. **G)** Structure of the I3 V_L_ highlighting hydrogen bonds formed by the N54 mutations. **H)** Aligned I5 and I4 V_H_ domains showing similar elbow angles. **I)** Map and fit for the I4 intermediate showing the region around residue R57. Map density around the R57 side chain is consistent with a closed V1 loop. **J)** Gaussian filtered I4 intermediate bound state map showing N156-glycan density. **K)** Map and fit for the I4 intermediate of the region around residue R84. Limited density for the R84 side chain precluded definition of its possible interactions. **L)** Alignment of the I5 and I4 structures with the I4 map showing slight rearrangement in the antibody loop adjacent to the N332 glycan.

**Supplemental Figure 8.**
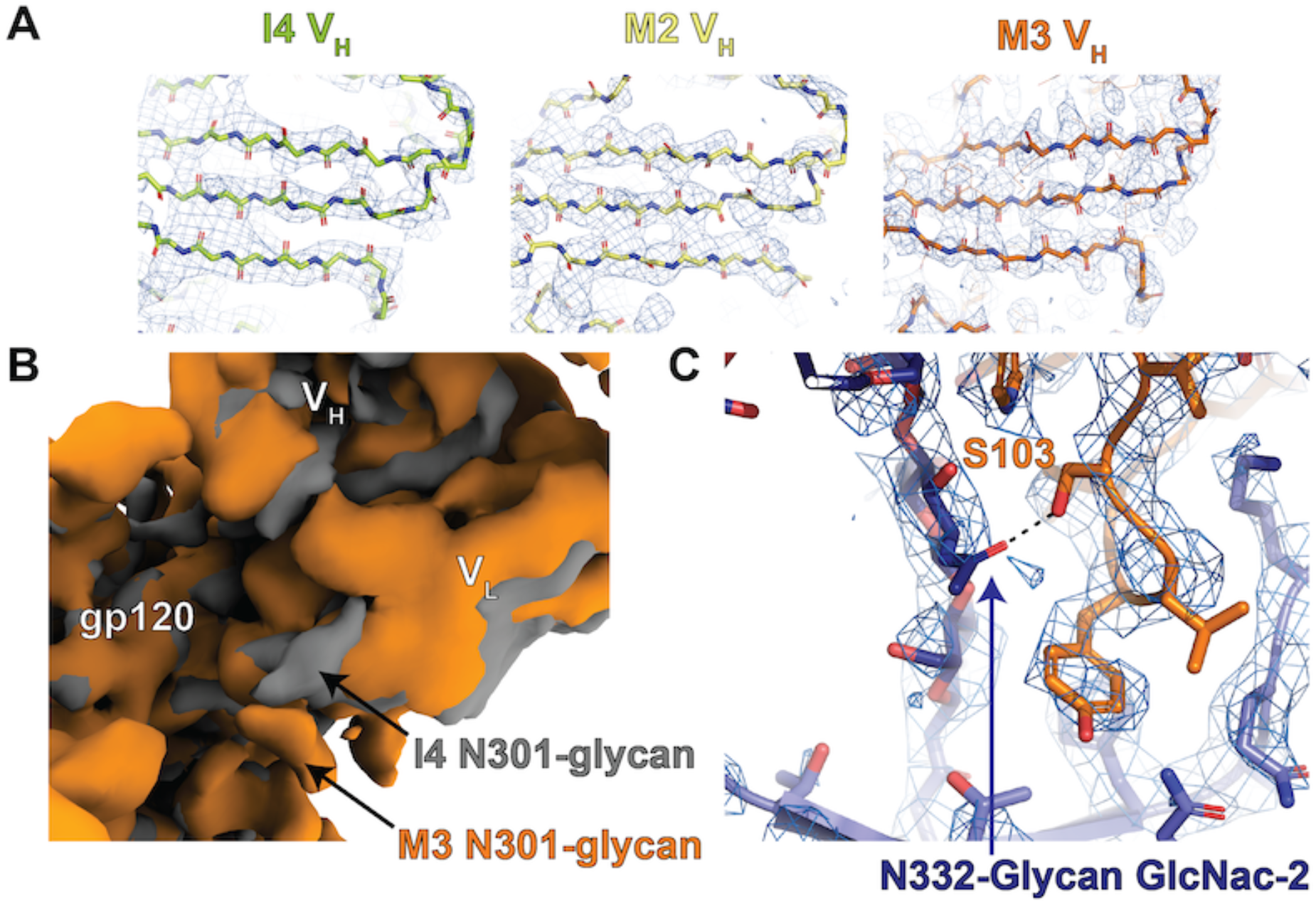
The I4 path split structures: Mature antibodies DH270.2 and DH270.3. **A)** Map and backbone structure fits for the I4, DH270.2 (M2) and DH270.3 (M3) heavy chains V_H_ domains. **B)** Gaussian filtered map alignments of the I4 and DH270.3 bound maps highlighting the shift in antibody and N301-glycan positions. **C)** Map and fit of the DH270.3 structure in the HCDR3 region highlighting hydrogen bonding between N332-glycan GlcNac-2 and heavy chain residue S103.

**Supplemental Figure 9.**
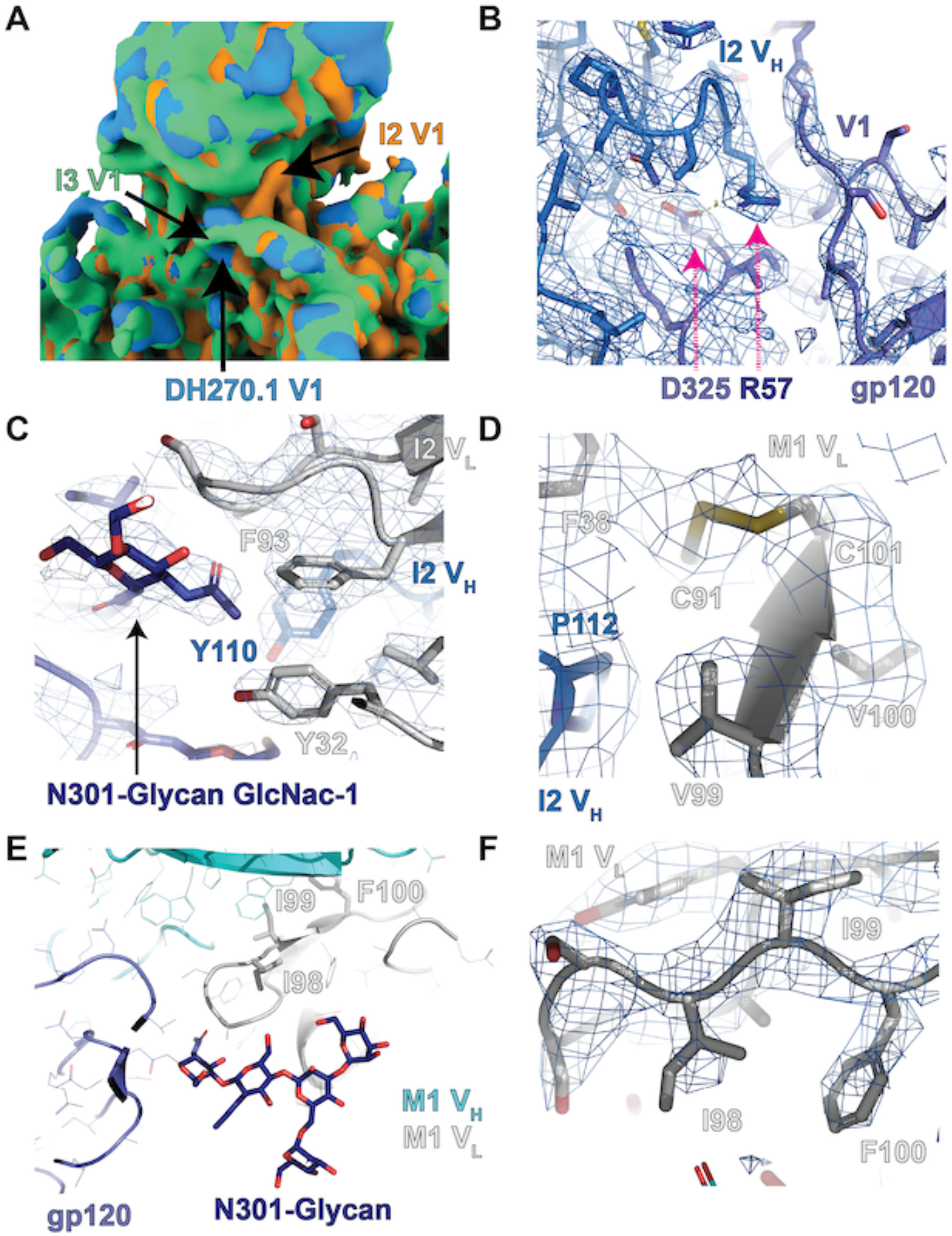
The I3 path split: Intermediate I2 and mature DH270.1. **A)** Gaussian filtered maps of the I3, I2, and DH270.1 (M1) bound maps showing differences in the V1 loop conformations. **B)** Map and fit of the I2 bound structure showing the shift in the I2 R57 side chain toward gp120 residue D325 of the GDIK motif. **C)** Map and fit of the I2 bound structure showing the position of Y110 relative to F93 and the N301-glycan base. **D)** Map and fit of the I2 bound structure showing the newly formed disulfide bond between C91 and the F101C site. **E)** DH270.1 structure near the LCDR3 mutations. **F)** Map and fit of the DH270.1 bound structures showing the position of the acquired hydrophobic mutations near LCDR3.

**Supplemental Figure 10.**
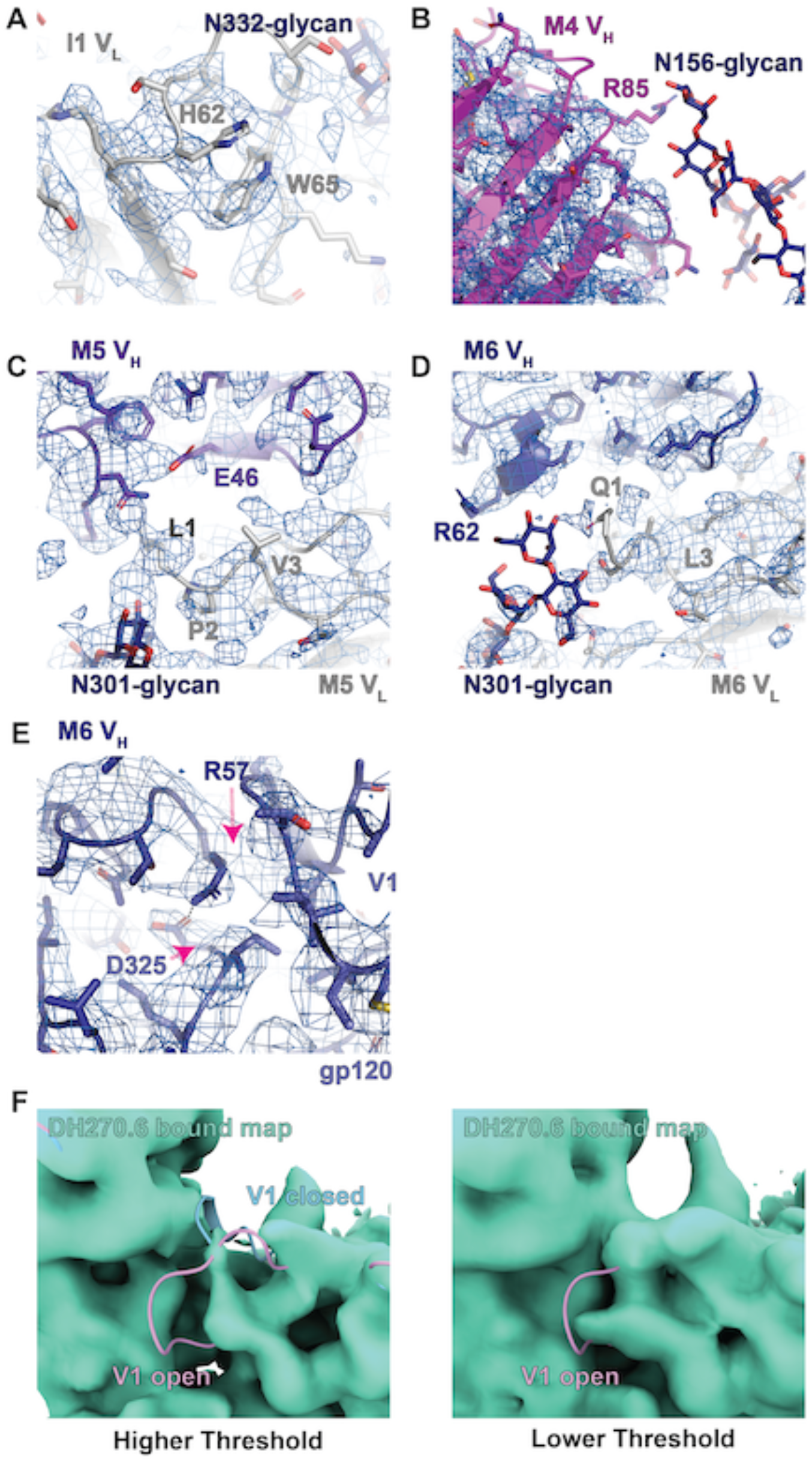
The I2 path split: Intermediate I1 and mature antibodies DH270.4, DH270.5, and DH270.6. **A)** Map and fit of the I1 bound structure at the N332-glycan adjacent light chain loop showing limited density for the mutation sites. **B)** Map and fit of the DH270.4 (M4) bound structure showing that poor map densities at R85 limit inspection of possible N156-glycan interactions. **C)** Map and fit of the DH270.5 (M5) bound structure in the region of the light chain N-terminus highlighting nearby mutations. **D)** Map and fit of the DH270.6 (M6) bound structure in the region of the light chain N-terminus highlighting nearby mutations. **E)** Map and fit of the DH270.6 (M6) bound structure showing the closed gp120 V1 loop and the shifted R57 side chain. **F)** Gaussian filtered map and closed and open V1 loops of the DH270.6 bound structure showing evidence of both a closed and an open state V1.

**Supplemental Figure 11.**
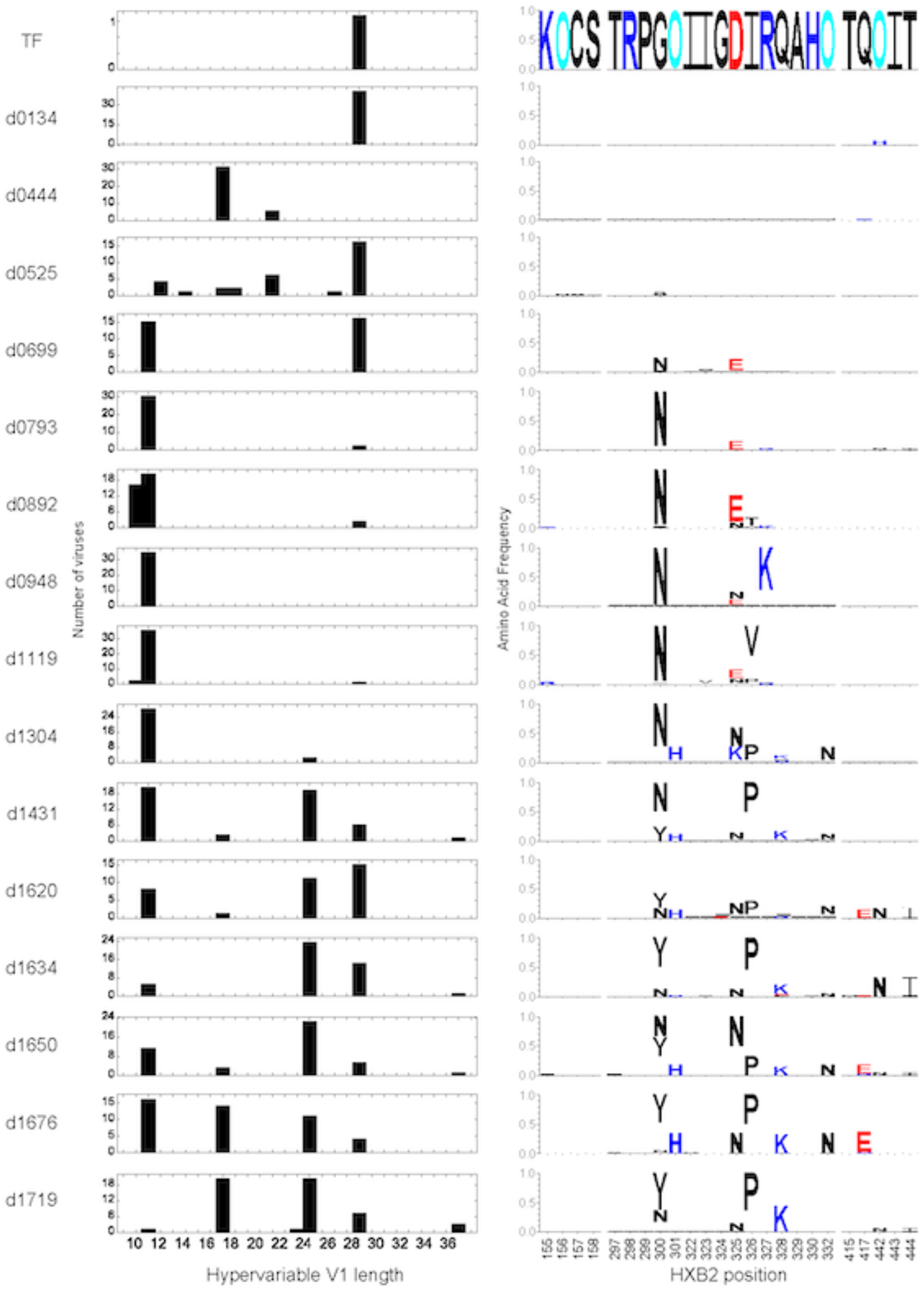
Longitudinal Env evolution in CH848 at key Env sites. The left panels show the distribution of hypervariable V1 loop lengths for longitudinally sampled CH848 Envs at each time point indicated to left of each plot (TF = transmitted founder; d0134 = 134 days post infection). The right panels show frequencies of amino acid and glycan mutations that arise over time at structurally key Env sites. For each plot, the height of the amino acid is proportional to its frequency at that time point, and the TF amino acid, shown only in the top LOGO, is blanked out to accentuate the mutations. Charge-based color-coding of amino acids is used (red: Asp, Glu; blue: His, Arg, Lys; black: all other amino acids), and potential N-linked glycan sites are indicated by “O” and colored cyan.

**Supplemental Figure 12.**
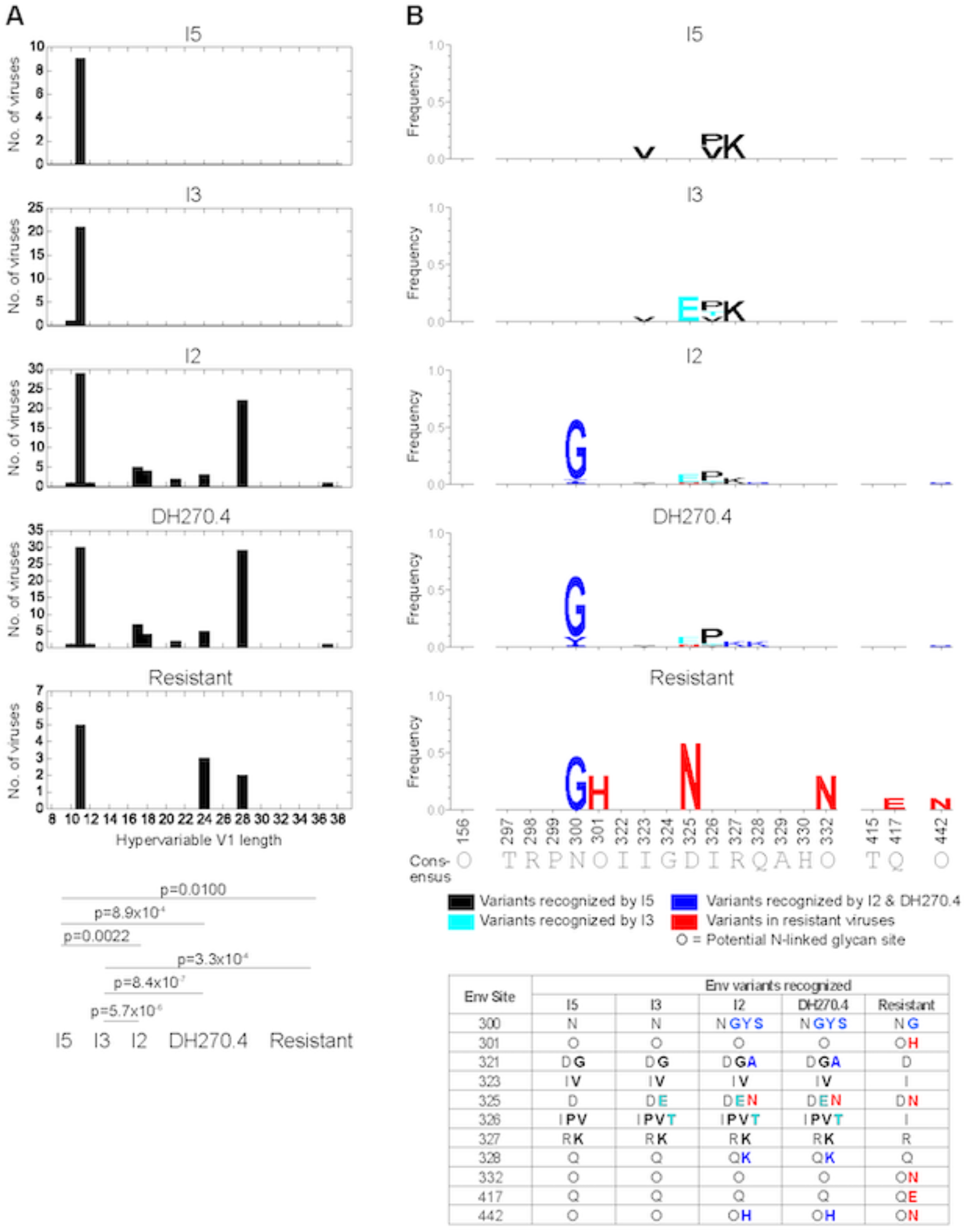
Env features of autologous viruses neutralized by intermediate and mature DH270 Abs. **A)** Histograms of hypervariable V1 loop lengths for the group of viruses that could be neutralized by I5, I3, I2 and DH270.4, arranged from top to bottom. The bottom-most plot shows histogram of hypervariable V1 loop lengths for autologous viruses that are resistant to all above antibodies. These V1 length distributions across antibodies were statistically compared using Wilcoxon Rank Sum Test and the bottom panel shows comparisons that yielded p-values < 0.05. **B)** Similar to panel (A) top, except amino acid frequency are shown using logos. To highlight diversity recognition, the amino acids and N-linked glycosylation sites of the consensus form are blanked out of the LOGOs; the consensus form is written along the bottom. The TF form matched the consensus except at position 300, which was Glycine. Variants are color-coded according to which intermediate could first recognize the variant, or if the variants were predominantly found in the resistant virus group. The table below summarizes the variants recognized by each antibody at the variable Env sites.

**Supplemental Figure 13.**
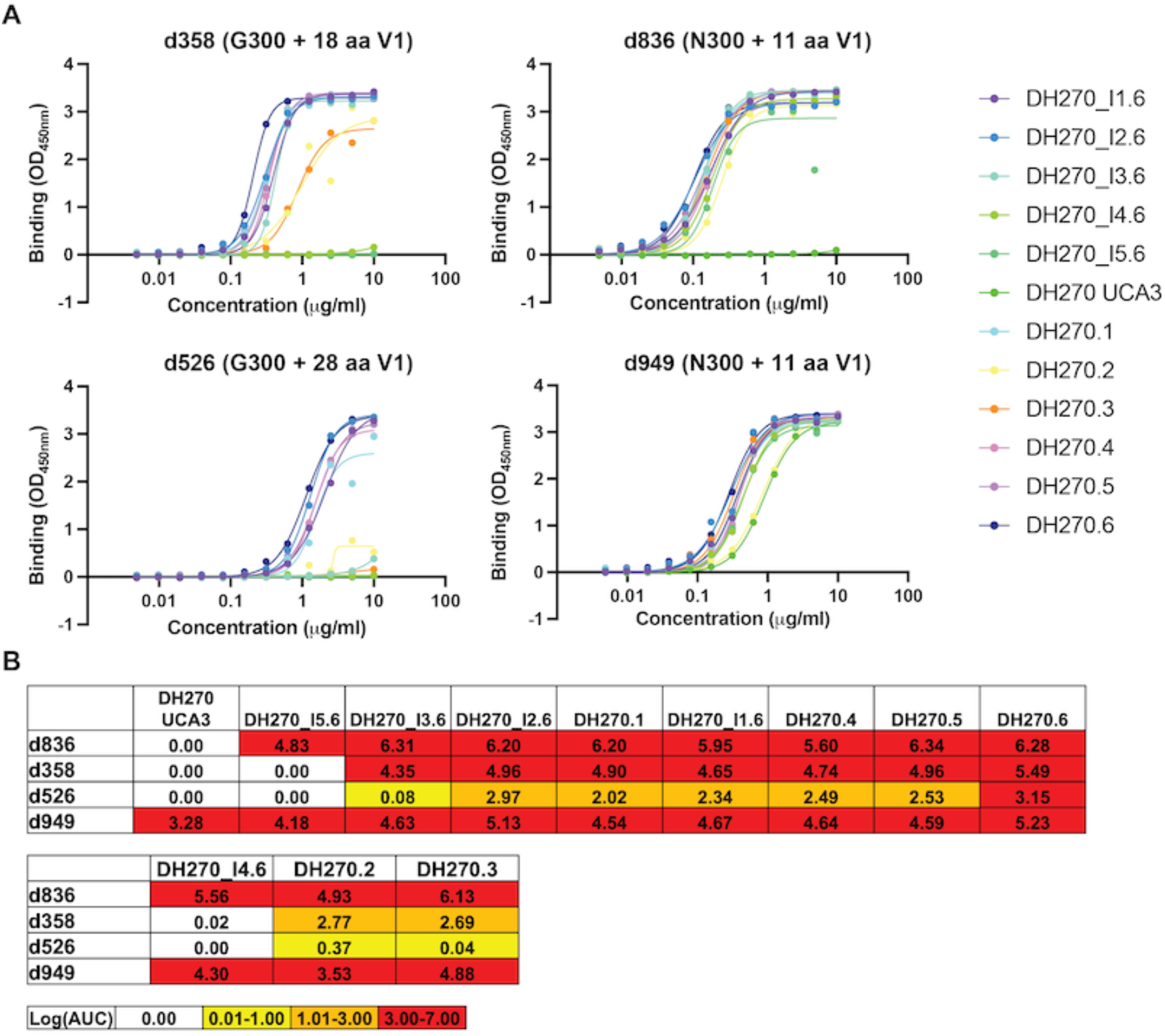
Binding of DH270 clonal clone antibodies to CH848 Env SOSIPs measured by ELISA. **A)** ELISA binding curves for CH848 d358, d526, d836, and d949. **B)** Table of log area under the curve for ELISA binding curves.

**Supplemental Figure 14.**
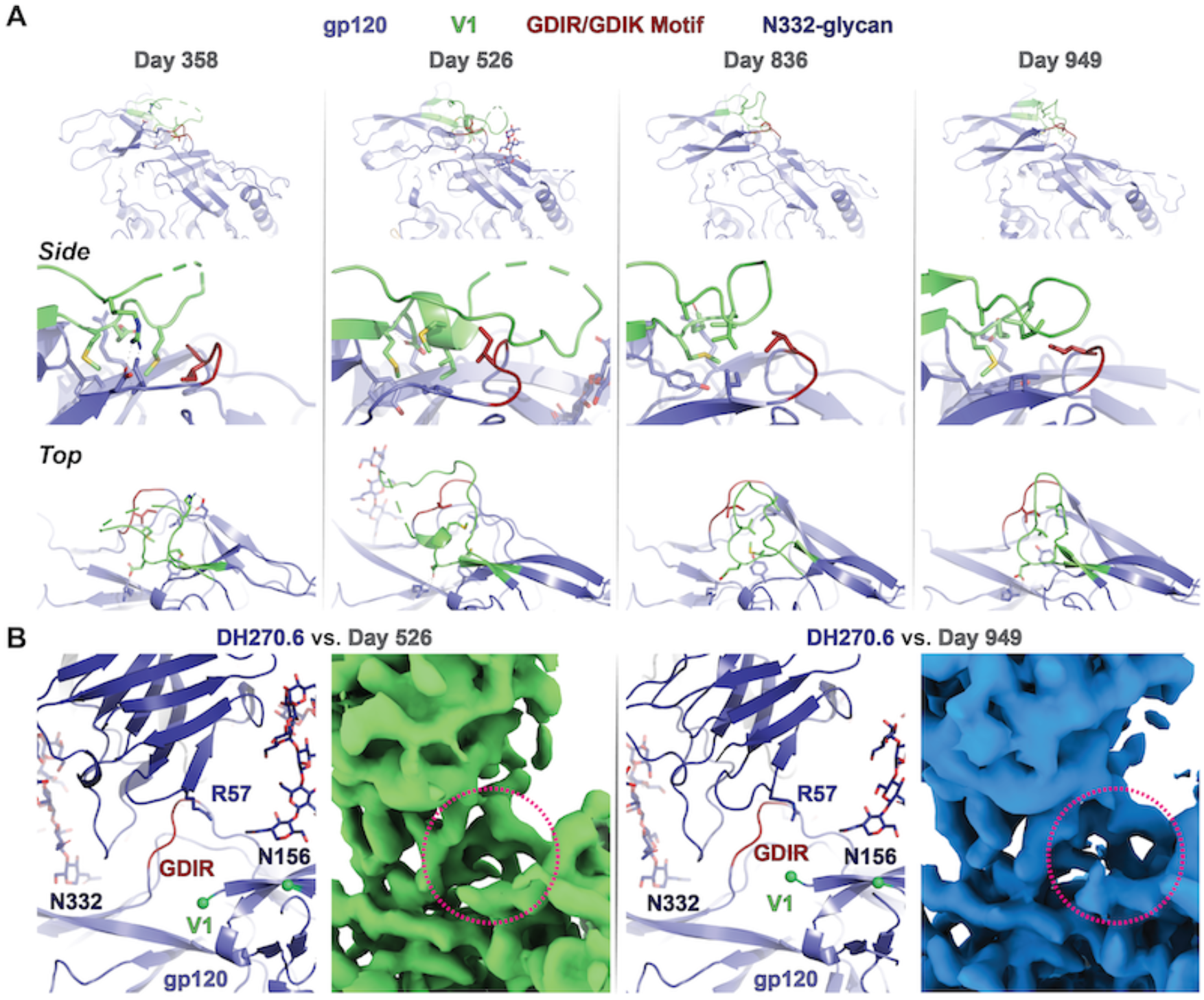
Variability in V1 loop length and sequence retains vulnerable Env motif protection. **A)** Structures of unliganded V3-glycan epitope Env ectodomains for Env sequences isolated from the CH848 infected individual at days 358, 526, 836, and 949 post infection. Upper panels depict a single Env gp120 (blue) protomer with the V1 loop (green) and the GDIR/K motif (red) highlighted. The middle panels present zoomed in, side views of the V3-glycan epitope showing the conserved hydrophobic core residues in stick representation. The lower panel presents a top view of the epitope. **B)** Comparison between the DH270.6 bound d526 and d949 trimer ectodomains showing the fit structure and maps. Green spheres in the structures identify the residue positions at which the gp120 V1 loops are poorly resolved in the maps. The magenta circle over the maps indicates the position of missing gp120 V1 loop densities.

**Supplemental Figure 15.**
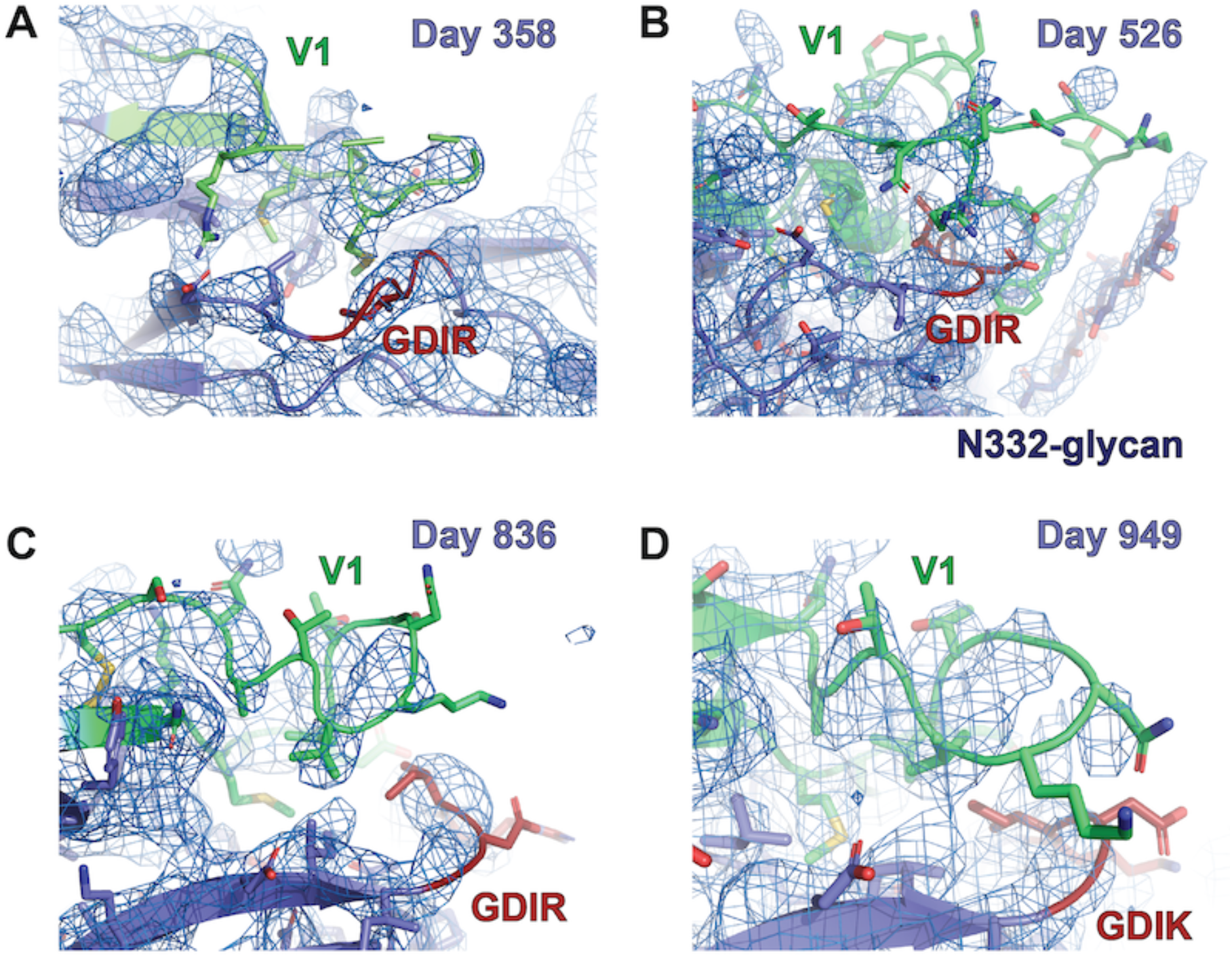
Unliganded V3-glycan epitopes of CH848 Env with different V1 loop lengths. **A-D)** Map and fits of the CH848 day 358, 526, 836, and 949 structures at the V3-glycan epitope.

## Acknowledgements

Cryo-EM data were collected at the Duke Krios at the Duke University Shared Materials Instrumentation Facility (SMIF), a member of the North Carolina Research Triangle Nanotechnology Network (RTNN), which is supported by the National Science Foundation (award number <u>ECCS-2025064</u>) as part of the National Nanotechnology Coordinated Infrastructure (NNCI), and at the National Center for Cryo-EM Access and Training (NCCAT) and the Simons Electron Microscopy Center located at the New York Structural Biology Center, supported by the NIH Common Fund Transformative High Resolution Cryo-Electron Microscopy program (<u>U24 GM129539</u>) and by grants from the Simons Foundation (<u>SF349247</u>) and New York State. This study used the computational resources offered by Duke Research Computing (http://rc.duke.edu; NIH 1S10OD018164-01) at Duke University. We thank Advaiti Kane and Parth Patel for antigenicity analysis of trimers.

## Author Contributions

R.H., A.B., and P.A. led the study; R.H., Y.Z., V.S., and P.A. determined and analyzed cryo-EM structures; V.S. expressed and purified proteins; K.O.S. supervised expression and purification of proteins; K.W. and B.K. performed and analyzed neutralization signatures; K.A. performed SPR assays; S.M.A. supervised SPR assays; M.B. and R.P. performed ELISA assays. R.H. wrote and edited the manuscript with help from all authors. B.F.H, and P.A. designed, edited, and acquired funding for the study. P.A. supervised the study and reviewed all data.

